# Formation of amyloid-like HTTex1 aggregates in neurons, downregulation of synaptic proteins and early mortality of Huntington’s disease flies are causally linked

**DOI:** 10.1101/2025.02.06.636778

**Authors:** Anne Ast, Leonard Roth, Lydia Brusendorf, Franziska Schindler, Orchid Wael Mostapha Ammar, Simon Berberich, Juliane Edel, Megan Bonsor, Elisabeth Georgii, Christian Hänig, Claudia Langnick, Andranik Ivanov, Dieter Beule, Marie Piraud, Severine Kunz, Oliver Popp, Philipp Mertins, Astrid G. Petzoldt, Erich E. Wanker

## Abstract

Amyloidogenic mutant huntingtin exon-1 (mHTTex1) protein aggregates with pathogenic polyglutamine (polyQ) tracts are the potential root cause of Huntington’s disease (HD). Here, we assessed the gain-of-function toxicity of mHTTex1 aggregation in neurons of HD transgenic flies. We show that the rate of mHTTex1 aggregation in neurons and early mortality of HD transgenic flies are correlated. We observed sequestration of key synaptic proteins into amyloid-like mHTTex1 aggregates and a concomitant decrease of their transcript levels, suggesting that progressive mHTTex1 aggregate stress in neurons leads to an impairment of synaptic function. Machine learning-based data analysis revealed that the abundance of synaptic proteins such as the vesicular monoamine transporter Vmat in the brain is predictive of fly survival. RNAi knockdown of Vmat-encoding transcripts in neurons with pathogenic amyloid-like HTTex1Q97 aggregates further shortened the lifespan of HD flies, supporting the hypothesis that mHTTex1 aggregation drives impairment of synaptic processes and pathogenesis of HD.

## INTRODUCTION

Huntington’s disease (HD) is a rare monogenic neurodegenerative disorder, manifesting with a triad of motor, psychiatric and cognitive symptoms, ultimately leading to premature death ^1^. HD is caused by the abnormal expansion of an unstable CAG repeat in the *HTT* gene. A greater CAG repeat number in exon 1 of the *HTT* gene correlates with an earlier age of onset and faster disease progression ^2,3^. Expression of *HTT* with a CAG repeat expansion leads to the production of a mutant huntingtin (mHTT) protein with an N-terminally elongated polyglutamine (polyQ) tract ^4^. Truncated N-terminal HTT fragments with abnormally expanded polyQ tracts have a high propensity to misfold and self-assemble into insoluble amyloidogenic protein aggregates with a fibrillar morphology in cell-free assays and *in vivo* model systems ^5,6^. In patient brains, N-terminal mHTT fragments with pathogenic polyQ tracts accumulate in neuronal inclusion bodies (IBs), which are a pathological hallmark of HD ^7^. Despite the genetic cause of HD having been known for more than 25 years, the pathological mechanisms underlying selective neurodegeneration and the development of symptoms remain largely unclear.

Understanding HD pathogenesis requires the identification and characterization of the molecular determinants that drive cellular dysfunction and neurodegeneration. Various potentially pathological RNA and protein species have been identified in the brains of HD models and patients, such as mHTT transcripts with elongated CAG repeats, soluble mHTT oligomers with expanded polyQ tracts or seeding-competent mHTTex1 fibrils ^6^. Evidence was presented that mRNAs with expanded CAG repeats can induce toxicity in model systems ^8^, suggesting that *HTT*-encoded transcripts might be causative to disease, independently of protein translation. mRNA transcripts with expanded CAG repeats were shown to form hairpins and sequester proteins into nuclear RNA foci, implying that abnormal mRNA-protein interactions may influence gene expression and/or splicing in HD brains ^9,10^. Dysregulation of gene transcription and mis-splicing are characteristic features of HD development ^11^. However, it needs to be determined whether they are caused by CAG-expanded mHTT transcripts or rather by polyQ-expanded mHTT protein molecules.

Several studies have demonstrated that expression of full-length (FL) polyQ-expanded mHTT in mouse brains is associated with HD-related molecular changes and symptoms ^12,13^. This suggests that soluble FL mHTT, readily detectable with biochemical methods in HD mouse brain extracts ^14^, is the neurotoxic protein species driving pathogenesis. However, small N-terminal polyQ-containing HTT fragments that are released from FL protein by proteolytic cleavage ^15,16^ or produced by other molecular mechanisms such as mis-splicing ^17,18^ are also always present in neuronal cells. These protein fragments have relatively low abundance and are often hard to detect with imaging and biochemical methods ^18^. Nevertheless, they may contribute substantially or even be responsible for the neurotoxicity and behavioural phenotypes observed in patients and HD models ^19^. Besides FL mHTT, various mHTT protein species including different types of aggregates (oligomers, protofibrils or fibrils) ^6,20^, which most likely have very different biological activities from soluble FL mHTT, may contribute to pathogenesis ^21^.

Our own recent studies revealed that small self-propagating, fibrillar mHTT assemblies formed of N-terminal fragments are prominent aggregate species in brain extracts of zQ175 knock-in HD mice ^22^. Importantly, these structures appear long before disease onset and progressively increase in abundance over time ^23^, suggesting that their accumulation in neurons may be responsible for the development of later symptoms ^24,25^. The disease-relevance of fibrillar mHTT aggregate species is also supported by studies indicating that they induce toxicity in cell models *per se* ^26,27^. Recent investigations with an inducible expression system in a transgenic HD *Drosophila* model have demonstrated that short-time production of small amounts of seeding-competent mHTTex1 fibrils with a pathogenic polyQ tract in adult neurons was sufficient to cause a dramatic shortening of the flies’ lifespan ^23^. This suggests that very low amounts of amyloidogenic mHTTex1 fibrils can induce a neurotoxic phenotype in HD flies. Besides fibrillar mHTTex1 species, prefibrillar protein assemblies such as small, diffusible mHTT oligomers ^28,29^ may also be critical to pathogenesis. Furthermore, polyQ-independent translational products synthesized by repeat-associated non-ATG-dependent (RAN) translation might induce toxicity in HD neurons ^30,31^. To this day, a variety of HD-specific protein and RNA molecules has been described ^6,32^ that may be relevant to pathogenesis either alone or in combination.

The causal chain of events by which potentially neurotoxic mHTT protein species cause motor impairment, alterations in synaptic function and plasticity or further phenotypes is still unknown. Time-dependent assessment of the transcriptome or the proteome in different mouse brain tissues has revealed a multitude of mHTT-associated molecular changes, suggesting that subcellular processes like the transport of membrane vesicles along microtubules, synaptogenesis or the release of mitochondrial RNA are impaired ^12,33^. However, a direct causal link between a defined polyQ-expanded mHTT protein species and the observed molecular changes could not be established from the currently available data. This is also the case when HTT-associated interactome data sets are considered ^34,35^. They suggest that HTT plays a functional role in diverse subcellular processes such as the regulation of gene transcription, formation and transport of autophagosomes or the control of protein translation ^4,6^. Information on whether alterations in interactions in these processes are linked to commonly observed disease phenotypes such as motor impairment or sleep deficits ^36^ remains scarce. More precise genotype-phenotype relationships need to be defined in relevant HD models to predict clinical manifestations that are indeed caused by a specific pathogenic mHTT protein species in neurons.

To establish genotype-phenotype relationships, we investigated the impact of mHTTex1 protein variants with different aggregation propensities on the development of molecular and behavioural phenotypes in HD transgenic flies. We systematically quantified mHTTex1 aggregation, sequestration of neuronal proteins into aggregates, global transcriptional dysregulation and survival of different HD fly strains under several experimental conditions. We found that the abundance of stable, amyloidogenic mHTTex1 aggregates in neurons and early mortality of HD transgenic flies are correlated, supporting previous observations that mutant HTTex1 aggregation and neurotoxicity are linked ^23^. Also, we observed that stable, pathogenic mHTTex1 aggregates sequester key synaptic proteins and concomitantly these proteins are transcriptionally downregulated in HD fly heads, suggesting that these phenomena are causally linked and contribute to the observed neurotoxicity and progressive development of disease-like phenotypes in HD transgenic flies with pathogenic mHTTex1 aggregates. This included the vesicular monoamine transporter Vmat, a key regulator of neurotransmission ^37^. A machine-learning-based analysis of both transcriptome and survival data sets of HD and control fly strains revealed that the abundance of synaptic proteins involved in neurotransmission is predictive of fly survival. This suggests that their transcriptional downregulation in fly neurons with amyloid-like HTTex1 aggregates contributes to terminal phenotypes such as motor impairment or premature mortality. Importantly, the knockdown of Vmat-encoding transcripts with siRNA in neurons with HTTex1Q97 aggregates further shortened the lifespan of HD transgenic flies, supporting the importance of efficient neurotransmission for fly survival. The relationship between progressive mHTTex1 aggregation in neurons, sequestration of synaptic proteins and their transcriptional downregulation in brains of HD transgenic flies are discussed.

## RESULTS

### Formation of SDS-stable HTTex1Q97 aggregates in neurons is associated with early mortality of HD transgenic flies and impairment of locomotor activity

To assess the relationship between polyQ-mediated mHTTex1 aggregation and neurotoxicity *in vivo*, we generated transgenic fly strains that produce human non-pathogenic (HTTex1Q17) or pathogenic (HTTex1Q49 and HTTex1Q97) HTTex1 proteins in neurons using the established Elav-GAL4 driver system ^38^. In addition, control flies that pan-neuronally express an HTTex1 protein without a polyQ tract (HTTex1Q0) were generated (**Fig. 1a**). In all transgenic fly strains, the cDNAs encoding HTTex1 proteins were integrated into the same chromosomal locus using the PhiC31 integration system ^39^ to avoid chromosomal position effects. Analysis of head extracts prepared from 5-day-old transgenic flies confirmed that transcripts encoding HTTex1 proteins were produced in neurons of all strains (**Fig. 1b**). Analysis of the head extracts by SDS-PAGE and immunoblotting revealed no or very low levels of soluble HTTex1 proteins while a larger N-terminal HTT protein with a non-pathogenic polyQ tract (HTT513Q17) and a folded α-helical N-HEAT domain (**Fig. 1c**, **Supplementary** Fig. 1a) was readily detectable in a control strain. High-molecular-weight HTTex1Q97 aggregates, migrating as a smear in gels, were clearly detectable (**Fig. 1c**), demonstrating that HTTex1 fragments with a long pathogenic polyQ tract rapidly self-assemble into stable, amyloid-like HTTex1 aggregates in young HD fly neurons. The formation of SDS-stable HTTex1Q97 aggregates in neurons of 3-day-old HD flies was confirmed using a well-established filter retardation assay (FRA) that exclusively detects large, SDS-and heat-stable amyloidogenic HTTex1 aggregates (**Fig. 1d**) but not small monomers or small unstable oligomers ^40^. Interestingly, amyloid-like HTTex1Q49 aggregates were also identified in head extracts of 50-day-old HD flies using a dot blot (DB) assay and the MW8 antibody, which preferentially detects HTTex1 aggregates ^41^. This indicates that a protein with 49Qs can form aggregates in neurons but at a much slower rate. In contrast, no HTTex1Q0 or HTTex1Q17 aggregates were identified with DBs in 50-day-old HD transgenic fly head extracts (**Fig. 1d**).

**Figure 1:**
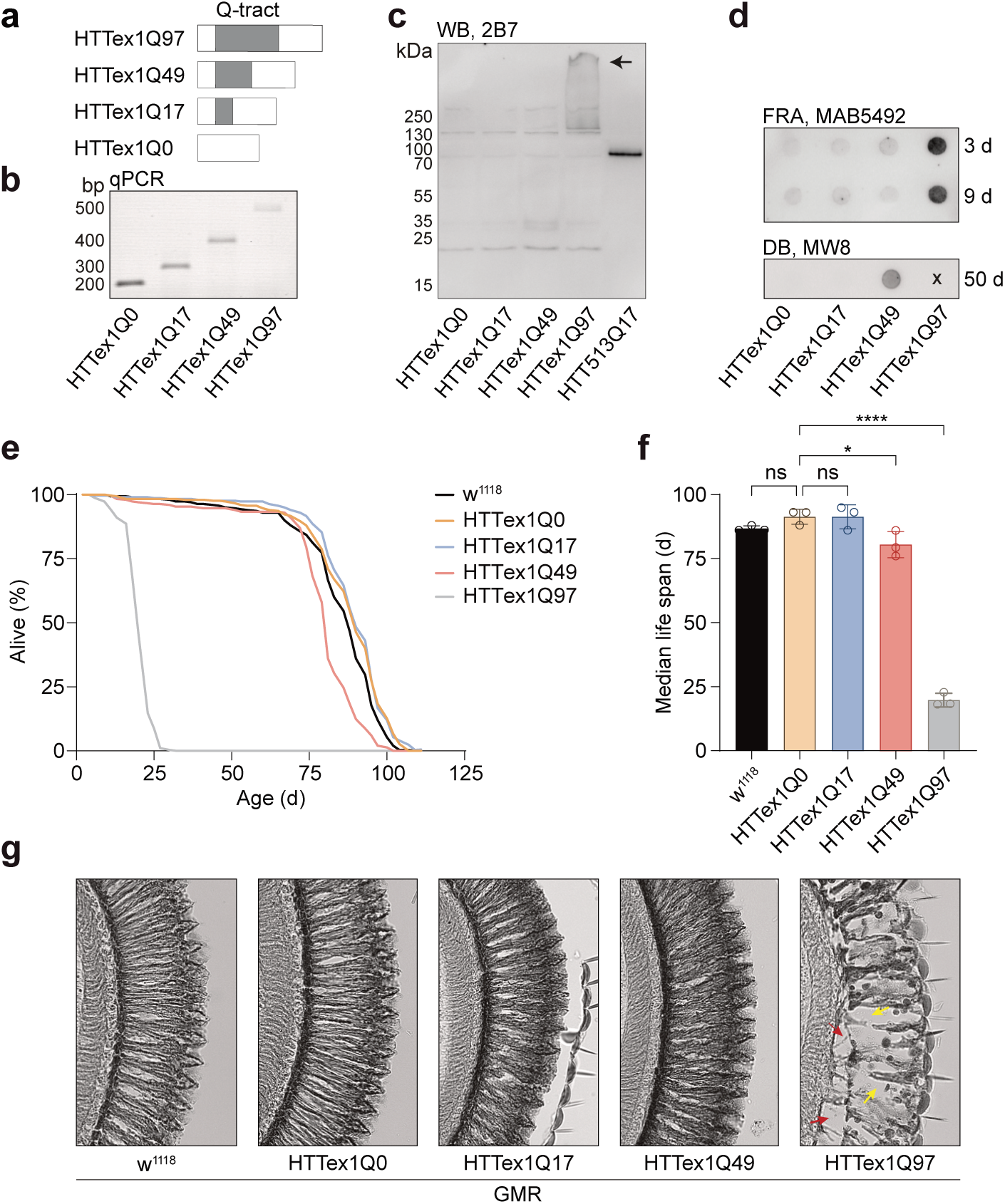
Formation of amyloid-like mHTTex1 aggregates in neurons is associated with early mortality of HD transgenic flies. **(a)** Schematic representation of HTTex1 fragments with and without polyQ tracts. The grey box represents the polyQ tract; the white boxes represent N- and C-terminal regions in the HTTex1 protein. **(b)** Analysis of HTTex1 transcripts in fly head extracts with qPCR; samples were prepared from Elav-GAL4-driven HD transgenic flies. Fragments with the expected sizes of ∼200 (HTTex1Q0), ∼250 (HTTex1Q17), ∼350 (HTTex1Q49) and ∼500 bp (HTTex1Q97) were amplified. **(c)** Analysis of head extracts prepared from HD transgenic flies pan-neuronally producing HTTex1 fragments or HTT513Q17 with the Elav-GAL4 driver by SDS-PAGE and immunoblotting. ∼75 µg total protein of HTTex1 fly lines was loaded onto the gel, while ∼7.5 µg total protein of the HTT513Q17 fly line was loaded. The membrane was probed with the anti-HTT antibody 2B7. The arrow indicates high molecular weight SDS-stable HTTex1 aggregates in the stacking gel. **(d)** Analysis of head extracts prepared from HD transgenic flies neuronally producing HTTex1 fragments with filter retardation (FRA) and dot blot (DB) assays. FRAs were performed with head extracts of 3- and 9-day-old flies; membranes were probed with the anti-HTT antibody MAB5492. DB assays with head extracts of 50-day-old flies. 15 µg of total protein was loaded and probed with the anti-HTT antibody MW8. **(e)** Survival analysis of Elav;HTTex1 transgenic flies; survival was plotted as the percentage of surviving flies of three biological replicates (each with n =∼100 flies/group). **(f)** Median life span calculated from survival curves **(e)**. Bars are mean ± SEM from 3 independent replicates; statistical significance assessed by One-way ANOVA Dunnett’s post-hoc test compared to HTTex1Q17; n.s., not statistically significant. **(g)** Impact of GMR-driven HTTex1 expression on photoreceptor degeneration. Shown are representative horizontal retina paraffin sections of 3-day-old female flies. As a negative control GMR;w^1118^ flies were analyzed. For each genotype ∼20 fly heads were analyzed. Yellow arrows indicate loss of retinal integrity and red arrows show the detachment of retinal floor and photoreceptor cells.

Next, we performed survival experiments with the different fly strains to assess the relationship between HTTex1 aggregation and neurotoxicity. We found that Elav;HTTex1Q0, Elav;HTTex1Q17 and Elav;w^1118^ control flies have a median lifespan of ∼85-90 days (**Fig. 1e and 1f**). In comparison, the survival of Elav;HTTex1Q97 transgenic flies, in which amyloid-like HTTex1Q97 aggregates were detectable in neurons already at a young age (**Fig. 1c and 1d**), was very short (median lifespan ∼20 days). Interestingly, a significantly shortened median lifespan (∼81 days) was also observed with Elav;HTTex1Q49 flies (**Fig. 1e and 1f**). In these flies, amyloid-like HTTex1Q49 aggregates accumulate at a slower rate compared to Elav;HTTex1Q97 flies (**Fig. 1c and 1d**). Together, these studies suggest that the formation of amyloid-like mHTTex1 aggregates, which occurs early in neurons of young flies, and the premature mortality of HD transgenic flies are causally linked.

The association of polyQ-dependent mHTTex1 aggregation in neurons and neurotoxicity was confirmed by quantification of the circadian locomotor activity ^42^ of HD flies using a *Drosophila* activity monitor (**Supplementary** Fig. 1b). We found that locomotor activity of 5-day-old Elav;HTTex1Q97 flies is significantly decreased in comparison to Elav;HTTex1Q0, Elav;HTTex1Q17 or Elav;HTTex1Q49 transgenic flies (**Supplementary** Fig. 1c and 1d). Finally, transgenic HD flies were crossed with GMR-GAL4 driver flies ^43^ to investigate the impact of pathogenic and non-pathogenic HTTex1 proteins on photoreceptor physiology. As expected, our analysis of retinal sections of 3-day-old GMR;HTTex1Q97 flies revealed dramatically damaged photoreceptor cells (**Fig. 1g**), while no such changes were found in young GMR;HTTex1Q0, GMR;HTTex1Q17 or GMR;HTTex1Q49 flies. Together these investigations support a causal relationship between mHTTex1 aggregation and the premature death of HD flies, which is only observed after stable, amyloid-like mHTTex1 aggregates are detectable in neurons of HD flies.

### Interruption of pathogenic polyQ tracts reduces aggregation and neurotoxicity of HTTex1 proteins in transgenic flies

Previous investigations have shown that interruptions in pathogenic polyQ sequences of mHTTex1 fragments decrease their propensity for spontaneous aggregation ^44^. Interruptions may also lead to reduced stability of amyloid fibrils. We hypothesized that HTTex1 proteins with interrupted pathogenic polyQ tracts may be less aggregation-prone and also less neurotoxic than HTTex1 proteins with uninterrupted pathogenic polyQ tracts. To address this question, we generated HD transgenic flies that pan-neuronally express mHTTex1 fragments with (Ex1Q61ENK14 and Ex1Q71P4) and without polyQ interruptions (Ex1Q75) using the established Elav-GAL4 driver system (**Fig. 2a**). We exchanged 10 Qs in the polyQ tract of Ex1Q75 with positively and negatively charged amino acids (5 lysines, K; 5 glutamic acids, E) to obtain protein variant Ex1Q61ENK14. Charged amino acids were previously shown to function as β-sheet breakers ^44^, suggesting that Ex1Q61ENK14 might have a lower aggregation propensity than Ex1Q75. In addition, four Qs were exchanged for asparagines (N) (**Fig. 2a; Supplementary Table 1**). These polar residues have biochemical properties similar to Qs. Because they are shorter than Qs, however they might reduce the formation of inter-chain hydrogen bonds, when amyloid-like HTTex1 fibrils are formed in neurons ^45^. Thus, we assumed amyloid-like Ex1Q61ENK14 fibrils formed *in vivo* to be less stable than Ex1Q75 fibrils. In the protein variant Ex1Q71P4 (**Fig. 2a**), the polyQ tract was interrupted with four prolines (P), which also function as β-sheet breakers ^46^. Therefore, Ex1Q71P4 might also aggregate less efficiently and form aggregates with reduced stability in transgenic HD flies than Ex1Q75. A transgenic fly strain pan-neuronally expressing a non-pathogenic Ex1Q17 protein was also generated (**Fig. 2a**). In contrast to HTTex1Q0, HTTex1Q17, HTTex1Q49 or HTTex1Q97 (**Fig. 1a**), these four HTTex1 proteins (Ex1Q17, Ex1Q61ENK14, Ex1Q71P4 and Ex1Q75) were C-terminally tagged with a short V5 epitope tag for immunodetection (**Fig. 2a; Supplementary Table 1**).

**Figure 2:**
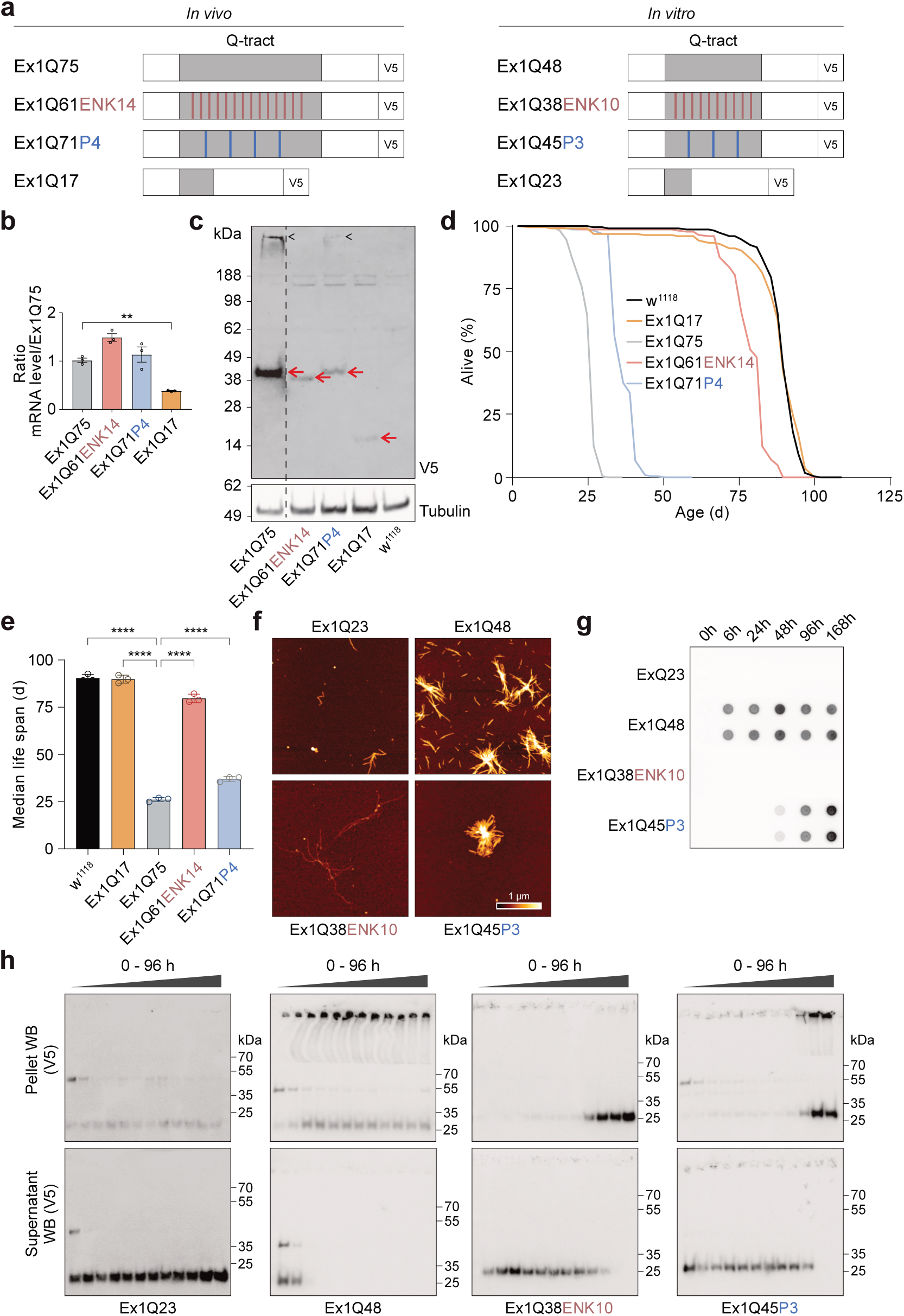
Interruptions of pathogenic polyQ tracts with charged amino acids or proline residues reduce mHTTex1 aggregation and neurotoxicity in HD flies. **(a)** Schematic representation of C-terminally V5-tagged HTTex1 protein variants (Ex1) with and without polyQ interruptions. Red indicates the exchange of Qs against glutamic acid (E), asparagine (N) and lysine (K) residues; blue indicates the exchange of Qs against prolines (P). V5 indicates the C-terminal V5-tag fused to the HTTex1 fragments for immunodetection. **(b)** Quantification of relative HTTex1 mRNA levels in head extracts of 10-day-old Elav-driven HD transgenic flies by qPCR; three biological replicates are shown. **(c)** Analysis of head lysates prepared from 10-day-old Elav-driven HD transgenic flies by SDS-PAGE and immunoblotting using the V5 antibody. 7.5 mg total protein was loaded per lane. **(d)** Lifespan analysis of Elav-driven HD transgenic flies expressing V5-tagged HTTex1 protein variants in neurons. Life span plotted as the percentage of surviving flies of 3 biological replicates (n_Ex1Q75_ = 86, 94, 88; n_Ex1Q61ENK14_ = 88, 83, 86; n_Ex1Q71P4_= 98, 95, 85; n_Ex1Q17_ = 93, 86, 90; n_w1118_ = 100, 87, 89). **(e)** Median life span calculated from survival curves in **(d)**. Average survival in each experiment is presented as white dots. Bars are mean ± SEM from 3 independent replicates; statistical significance assessed by One-way ANOVA Dunnett’s post-hoc test; data were compared to Ex1Q75 transgenic flies. **(f)** Atomic force microscopy (AFM) analysis of spontaneously formed HTTex1 protein aggregates. GST-HTTex1 fusions (**Supplementary** Fig. 2e) with and without polyQ interruptions (4 µM) were incubated with Prescission protease (PSP) and after 168 h samples were analyzed. Scale bar: 1 µm; color gradient represents 0-20 nm height. **(g)** Analysis of time-dependent aggregation of HTTex1 protein variants with and without polyQ interruptions using a filter retardation assay (FRA). GST-HTTex1 fusion proteins (4 µM) were incubated at 25°C with PreScission protease (PSP) for indicated times. 500 ng protein of each reaction were filtered through a cellulose acetate membrane. SDS- and heat-stable HTTex1 aggregates retained on filter membrane detected with an anti-V5 antibody. **(h)** Time-resolved analysis of spontaneous HTTex1 aggregation using a centrifugation assay followed by SDS-PAGE and immunoblotting. HTTex1 protein variants with and without polyQ interruptions were analyzed. After incubation of GST-HTTex1 fusions with PSP for 0, 1, 2, 3, 4, 5, 6, 7, 8, 24, 48 or 96 h samples were centrifuged (186,000 x g) to separate high (pellet) and low molecular weight HTTex1 protein species (supernatant) under non-denaturing conditions. Finally, pellet (top) and supernatant (bottom) fractions were analyzed by SDS-PAGE (denaturing conditions) and immunoblotting using the V5 antibody.

First, *HTTex1* transcript levels were quantified in heads of 10-day-old flies using an established qPCR method. We measured comparable *Ex1Q75*, *Ex1Q61ENK14* and *Ex1Q71P4* transcript levels in head extracts of HD transgenic flies (**Fig. 2b**). *Ex1Q17* transcript levels were significantly lower, suggesting that this protein is produced less efficiently in neurons than the pathogenic proteins Ex1Q61ENK14, Ex1Q71P4 or Ex1Q75. However, all four HTTex1 proteins were detected when head lysates were analysed by PAGE and immunoblotting (**Fig. 2c**). Interestingly, a higher abundance of Ex1Q75 than of Ex1Q61ENK14, Ex1Q71P4 or Ex1Q17 was observed. This suggests that the HTTex1 proteins with polyQ interruptions (Ex1Q61ENK14 and Ex1Q71P4) are degraded more efficiently in neurons than Ex1Q75, which contains an uninterrupted, pathogenic polyQ tract. Supporting this result, high-molecular-weight, SDS-stable aggregates were readily detectable in head extracts of Ex1Q75-expressing flies but not in those of Ex1Q61ENK14-, or Ex1Q17-expressing flies (**Fig. 2c**). A small amount of SDS-stable, high-molecular-weight aggregates was detectable also in head extracts of 10-day-old Elav;Ex1Q71P4 flies, indicating that this protein similar to Ex1Q75 forms very stable amyloid-like HTTex1 aggregates in neurons but with a much lower rate. An analysis of head extracts with FRAs confirmed the formation of SDS- and heat-stable Ex1Q75 and Ex1Q71P4 protein aggregates in fly neurons (**Supplementary** Fig. 2a and 2b), while such structures were undetectable in head extracts of Elav;Ex1Q61ENK14 or Elav;Ex1Q17 transgenic flies.

Finally, we compared the impact of HTTex1 aggregation in neurons on the survival of HD transgenic flies with and without interrupted polyQ tracts. Survival was measured over time by counting dead flies and by calculating median life spans (**Fig. 2d and 2e**). While Elav;Ex1Q17 flies had a normal median life span of ∼90 days, HD flies that rapidly accumulate SDS-stable Ex1Q75 in neurons had a median life span of ∼26 days. The median life spans of Elav;Ex1Q61ENK14 (∼80 days) and Elav;Ex1Q71P4 (∼37 days) flies were significantly extended, indicating that polyQ interruptions significantly prolong the survival of HD flies. Interestingly, the median life span of Ex1Q61ENK14 flies was significantly greater than of Ex1Q71P4 flies (**Fig. 2e**), suggesting that SDS- and heat-stable Ex1Q71P4 aggregates (**Supplementary** Fig. 2a and 2b) are more neurotoxic than Ex1Q61ENK14 aggregates, which do not possess such biochemical properties. Very similar results were obtained when HD transgenic flies pan-neuronally producing the proteins Ex1Q75, Ex1Q61ENK14, Ex1Q71P4 or Ex1Q17 with the Elav-GAL4 driver were assessed for locomotor activity using an established climbing assay (**Supplementary** Fig. 2c and 2d). Together these experiments substantiate our hypothesis that the abundance of SDS-stable, amyloid-like mHTTex1 aggregates in neurons and premature mortality of HD transgenic flies are causally linked.

### Interruption of pathogenic polyQ tracts in HTTex1 proteins reduces aggregation propensity and aggregate stability *in vitro*

To corroborate our *in vivo* studies with cell-free assays, we produced purified GST-tagged HTTex1 proteins (**Supplementary** Fig. 2e and 2f) with polyQ interruptions (Ex1Q38ENK10 and Ex1Q45P3) recapitulating the Ex1Q61ENK14 and Ex1Q71P4 proteins in flies (**Fig. 2a**) and studied their spontaneous aggregation *in vitro* using an established polymerization assay ^47^. As controls, HTTex1 fusion proteins without polyQ interruptions (Ex1Q23 and Ex1Q48) were investigated (**Supplementary** Fig. 2e and 2f). We observed that all four recombinant proteins (Ex1Q23, Ex1Q48, Ex1Q38ENK10 and Ex1Q45P3) are able to self-assemble into amyloid-like, fibrillar aggregates in cell-free assays (**Fig. 2f**). However, spontaneous HTTex1 aggregation *in vitro* occurs at very different rates. While first fibrillar Ex1Q48 aggregates were already detectable after 6 h, formation of fibrillar Ex1Q45P3 or Ex1Q38ENK10 aggregates was observed only after ∼48-96h (**Fig. 2f and 2g**). Incubation for 168 h was required until even small amounts of fibrillar Ex1Q23 aggregates could be detected (**Fig. 2f**). Detection of SDS- and heat-stable amyloidogenic HTTex1 aggregates with FRAs was only possible with the proteins Ex1Q48 and Ex1Q45P3 (**Fig. 2g**), confirming the results obtained *in vivo* with the related proteins Ex1Q75 and Ex1Q71P4 (**Fig. 2c, Supplementary** Fig. 2a and 2b).

To separate small from large mHTTex1 aggregates under non-denaturing conditions and subsequently assess their stability, samples of polymerization reactions were first centrifuged at high speed to obtain pellet and supernatant fractions (**Supplementary** Fig. 2g). Then, these fractions were analyzed by SDS-PAGE and immunoblotting to assess the SDS- and heat-stability of the small and large mHTTex1 assemblies obtained by centrifugation. SDS- and heat-stable HTTex1 aggregates are retained in the stacking gel, while instable structures are dissociated under these conditions and detected as monomers in SDS gels (**Supplementary** Fig. 2g). Our analysis confirmed the time-dependent formation of large, SDS-instable amyloid-like Ex1Q38ENK10 aggregates after ∼24h, while a mixture of both SDS-stable and instable aggregates was detectable with the Ex1Q45P3 protein (**Fig. 2h**). In strong contrast, Ex1Q48 without polyQ interruptions exclusively formed large, insoluble, SDS-stable aggregates that were retained in the stacking gel. Together, these studies support the hypothesis that Ex1Q61ENK14 forms neurotoxic amyloid-like protein aggregates in HD fly brains over time (**Fig. 2d and 2e**). In comparison to aggregates formed by Ex1Q75 or Ex1Q71P4, these structures are SDS- and heat-instable and therefore cannot be detected with FRAs, which exclusively detects SDS- and heat-stable amyloid-like mHTTex1 aggregates formed *in vitro* or *in vivo* ^40^.

### Progressive accumulation of amyloid-like HTTex1Q97 aggregates in adult neurons leads to sequestration of synaptic proteins

Our previous investigations indicate that the induction of HTTex1Q97 aggregation in young adult neurons with the RU486-inducible Elav-GeneSwitch (GS) driver system is sufficient to significantly shorten the lifespan of HD transgenic flies ^23^. Here, we wanted to assess the relationship between the time-dependent accumulation of amyloid-like HTTex1Q97 aggregates in adult neurons and the survival of HD flies. We started RU486 treatment in 3-day-old flies and quantified the formation of SDS-stable HTTex1Q97 aggregates in fly head extracts using FRAs (**Fig. 3a**). In parallel, the survival of RU486-treated elavGS;HTTex1Q97 HD flies was monitored. As shown in **Fig. 3b**, SDS- and heat-stable HTTex1Q97 aggregates accumulate progressively in fly neurons until they reach saturation after ∼12 days of RU486 treatment. We observed a progressive accumulation of HTTex1Q97 aggregates over ∼25 days until the flies started to dye rapidly within a few days (**Fig. 3b**), indicating that neurons can cope with high amounts of insoluble pathogenic HTTex1 aggregates for weeks until the system collapses.

**Figure 3:**
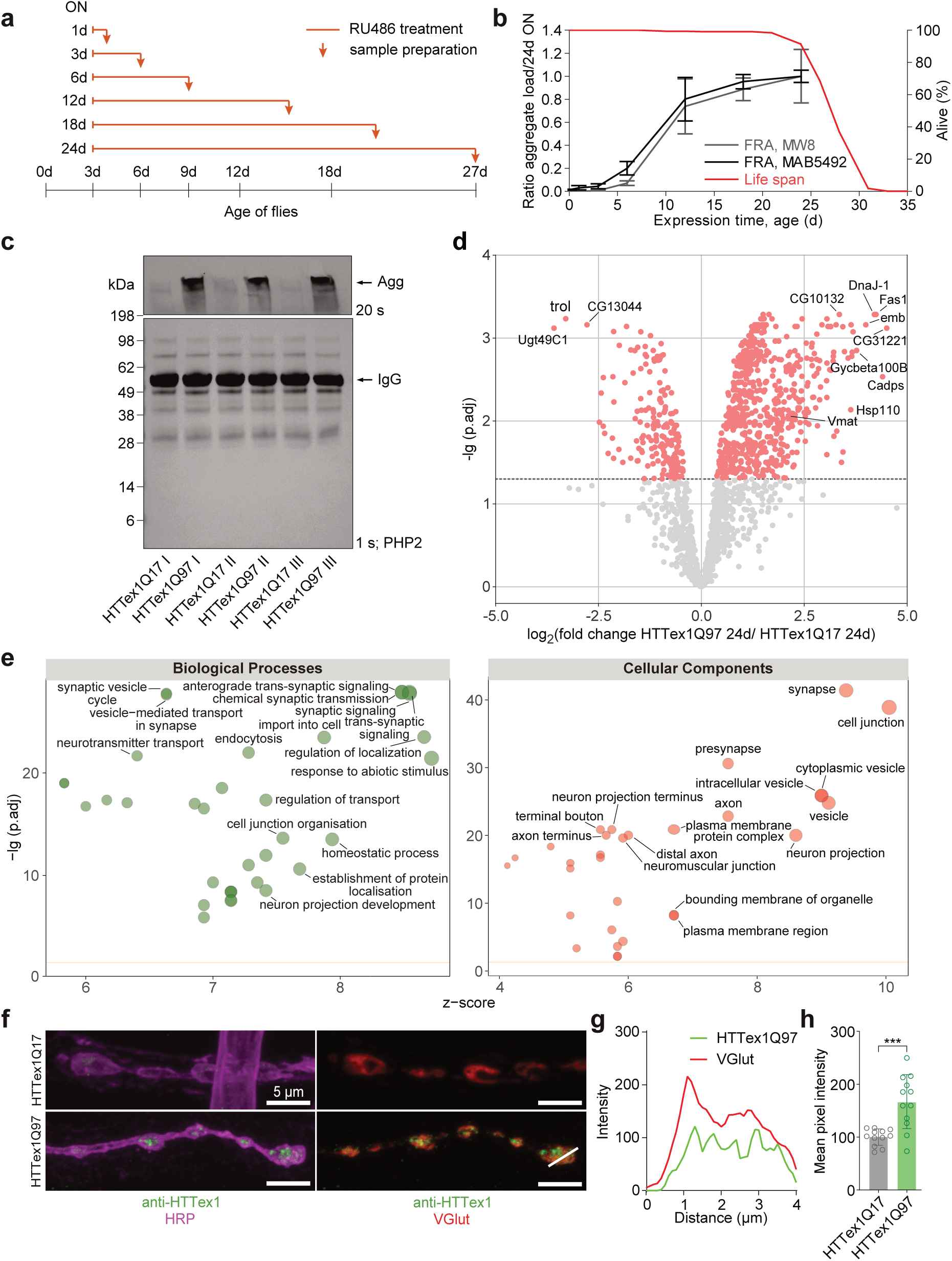
Progressive HTTex1Q97 aggregation in adult neurons leads to sequestration of synaptic proteins into amyloid-like structures. **(a)** RU486 treatment scheme to investigate the time-dependent accumulation of amyloid-like HTTex1Q97 aggregates in adult neurons of HD transgenic flies. Orange lines indicate the time periods of RU486 treatment; arrows indicate collection of fly heads for analysis. **(b)** Quantification of HTTex1Q97 aggregate load in fly head extracts and survival of elavGS;HTTex1Q97 flies after treatment of 3-day-old HD flies with the inducer RU486; see also scheme in (a). ∼15 µg of total protein prepared from fly heads at indicated time points were analyzed by filter retardation assay (FRA). Samples of multiple independent biological replicates (n = 3, 3, 3, 3, 3, 3, 2) were analyzed by FRA; error bars represent mean ± SEM. The red line displays the lifespan analysis of elavGS;HTTex1Q97 flies performed in three independent experiments (n = 141, 95, 99). MW8 and MAB5493 are anti-HTT antibodies. **(c)** Analysis of co-immunoprecipitates obtained from head extracts of 27-day-old elavGS;HTTex1Q97 and elavGS;HTTex1Q17 HD transgenic flies after induction of transgene expression with RU486 in 3-day-old flies (see also scheme in **(a)**. Samples were analyzed by immunoblotting and mass spectrometry (n = 3). Agg, high-molecular-weight SDS-stable HTTex1Q97 aggregates in the stacking gel; IgG, heavy chain; PHP2, anti-HTT antibody. **(d)** Volcano plot depicting the enrichment of *Drosophila* proteins with PHP2-immunoprecipitated HTTex1Q97 in comparison to PHP2-immunoprecipitated HTTex1Q17. Significantly enriched proteins (p.adj < 0.05) are indicated in red. **(e)** GO term enrichment analysis of proteins associated with HTTex1Q97 aggregates in dependence of normalized protein abundance (z-score). Significantly enriched GO terms (p.adj < 0.01, overlap >3, enrichment >1.5) are shown and a selection of terms associated with intracellular transport processes and synaptic functions are indicated. **(f)** Representative confocal immunofluorescence images of 3^rd^ instar *Drosophila* larvae of the neuromuscular junction (NMJ) expressing Elav;HTTex1Q17 (upper panel) or Elav;HTTex1Q97 (lower panel) labeled for HTT (green) or HRP (horseradish peroxidase, magenta) outlining the NMJ (left panel) and VGlut as synaptic vesicle marker (red, right panel). White line indicates line graph for **(g)**; scale bar 5 mm. **(g)** Line scan analysis supports the co-localization of HTTex1Q97 aggregates with VGlut at NMJs. **(h)** Quantification of the mean pixel intensities of the anti-HTT signal of (**f**) (n = 12; n represents NMJs) supports an accumulation of HTTex1Q97 aggregates in third instar larvae.

We hypothesized that the continuous sequestration of other neuronal proteins into amyloidogenic HTTex1Q97 aggregates might be responsible for the premature death of HD flies. To address this hypothesis, we treated 3-day-old animals with RU486 and then enriched amyloid-like HTTex1Q97 aggregates from head extracts of 27-day-old insects by co-immunoprecipitation (co-IP) using the monoclonal antibody PHP2, which preferentially binds amyloidogenic mHTTex1 aggregates ^48^. Antibody-enriched co-immunoprecipitates were analyzed by mass spectrometry (MS). A simplified workflow of this procedure is shown in **Supplementary** Fig. 3a. Control co-IP experiments with head extracts prepared from elavGS;HTTex1Q17 flies without amyloid-like mHTTex1 aggregates were also performed. As expected, enrichment of high-molecular-weight, SDS-stable HTTex1 aggregates was observed when immunoprecipitates of elavGS;HTTex1Q97 animals were analyzed by SDS-PAGE and immunoblotting (**Fig. 3c**), but not in immunoprecipitates of elavGS;HTTex1Q17 flies. Analysis of immunoprecipitates with MS revealed a significant co-enrichment of 618 proteins with HTTex1Q97 aggregates (**Fig. 3d and 3b; Supplementary Table 2a**). A gene ontology (GO) term enrichment analysis revealed that a large number of membrane-associated synaptic proteins such as Vmat (*Drosophila* vesicular monoamine transporter, ^49^, Brp (Bruchpilot, ^50^) or the RIM-binding protein RBP ^51^ were co-enriched with HTTex1Q97 aggregates (**Fig. 3e**), suggesting that progressive mHTTex1 aggregation in neurons influences critical synaptic processes. Analysis of co-IP samples with SDS-PAGE and immunoblotting confirmed the co-enrichment of Vmat and Brp with amyloid-like HTTex1Q97 aggregates (**Supplementary** Fig. 3c). Also, a co-localization of RBP with insoluble HTTex1Q97 aggregates was confirmed with immunohistology methods, when sections of whole HD fly brains were analyzed (**Supplementary** Fig. 3d and 3e).

Finally, we performed immunofluorescence-based co-localization studies in fixed *Drosophila* 3^rd^ instar larvae to assess whether HTTex1Q97 aggregates form and accumulate at the synaptic terminal, the neuromuscular junction (NMJ). Strikingly, our analysis revealed HTTex1Q97 aggregates at NMJs in Elav;HTTex1Q97 larvae, while such structures were undetectable in Elav;HTTex1Q17 larvae and Elav;w1118 driver controls (**Fig. 3f and Supplementary** Fig. 3f). We observed a drastic increase of mHTTex1 signal (mean pixel intensity over the entire NMJ) compared to controls (**Fig. 3g, 3h, Supplementary** Fig. 3g**, 3h**), supporting our hypothesis of an accumulation of pathogenic mHTTex1 assemblies at NMJs. Importantly, these experiments also revealed a partial co-localization of the synaptic glutamate transporter VGlut ^52^ and of Brp with amyloid-like HTTex1Q97 aggregates at NMJs (**Fig. 3f, 3g, Supplementary** Fig. 3g**, 3h**), confirming our co-IP MS data obtained with head extracts of adult HD flies. Brp is a well-characterized bona-fide presynaptic marker protein^50,53^.

### Formation of amyloid-like HTTex1Q97 aggregates in fly neurons is associated with transcriptional downregulation of synaptic proteins

Our studies with the RU486-inducible expression system indicate that progressive HTTex1Q97 aggregation in neurons leads to a sequestration of synaptic proteins into aggregates (**Fig. 3c-3h**) and that these abnormal protein-protein interactions might be the molecular cause for neuronal dysfunction and the observed premature mortality of HD flies. We hypothesized that HTTex1Q97 aggregation might also lead to transcriptome changes in brains of adult HD flies. To address this hypothesis, we expressed HTTex1Q97 or HTTex1Q17 for 3 or 6 days in 3-day-old transgenic flies by transferring them onto food with the hormone RU486 and then back to food lacking the inducer (3d-ON/OFF; 6d-ON/OFF; **Fig. 4a**). Previous studies have shown that short-time production of HTTex1Q97 aggregates in young adult neurons with RU486 is sufficient to induce dysfunction and premature fly mortality. In contrast, short-term neuronal expression of HTTex1Q17 does not lead to aggregation or influence survival ^23^. In addition, transgenic flies (elavGS;HTTex1Q97 and elavGS;HTTex1Q17) were continuously maintained on food with (ON) or without the inducer (OFF). mRNA was prepared from fly heads and quantitative transcriptome (RNAseq) data sets were produced with next generation sequencing after 27 days (**Fig. 4a**). At this time point many synaptic proteins are already sequestered into amyloid-like mHTTex1 aggregates in HD fly brains (**Fig. 3c-3e**).

**Figure 4:**
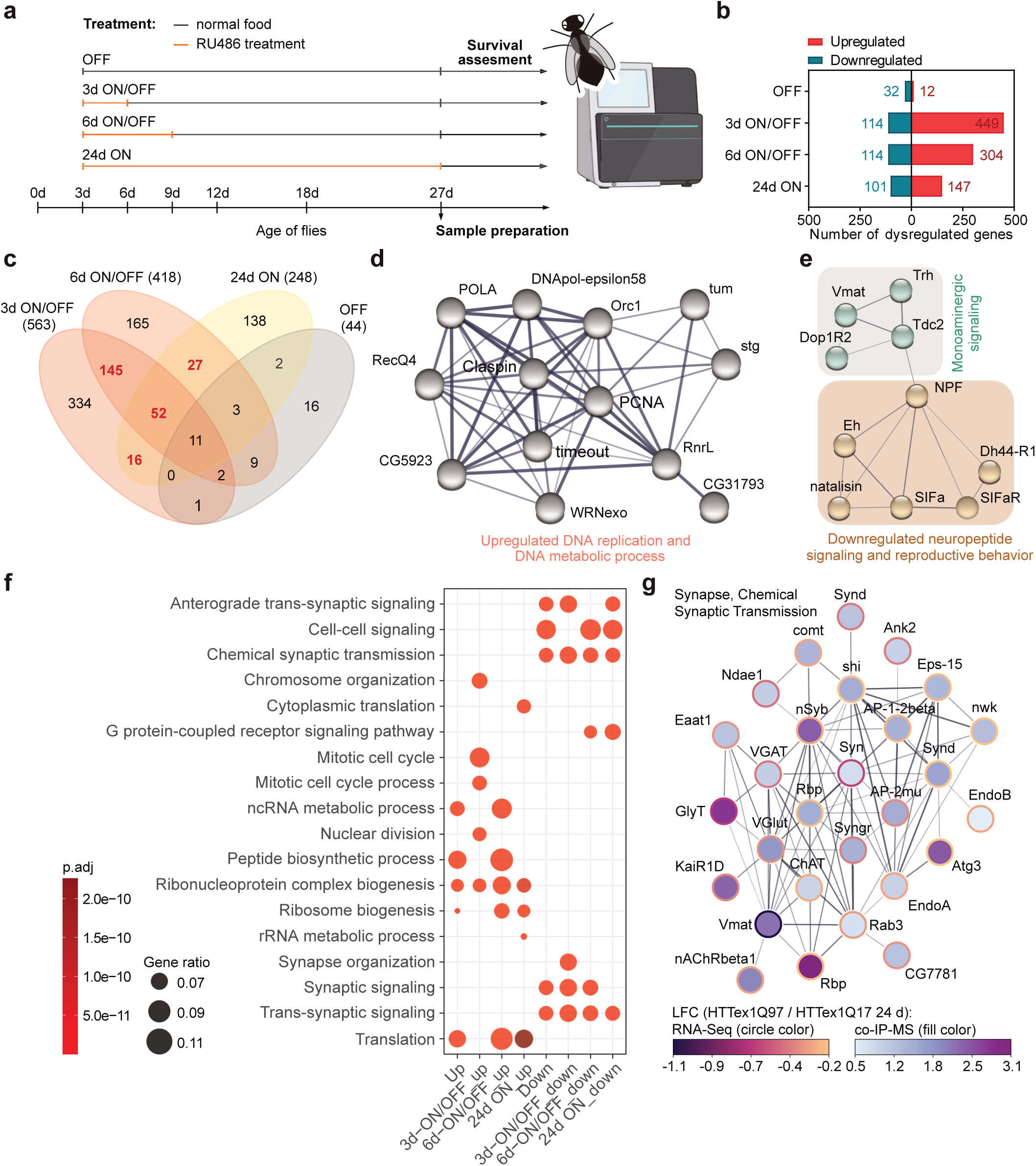
Accumulation of amyloid-like HTTex1Q97 aggregates in fly neurons is associated with downregulation of transcripts encoding synaptic proteins. **(a)** Scheme depicting short- and long-time RU486 treatment of adult elavGS;HTTex1Q97 or elavGS;HTTex1Q17 HD transgenic flies for transcriptome analysis. Fly head extracts of 27-day-old Q97 and Q17 (control) animals were utilized for mRNA preparation and RNA-seq analysis. **(b)** Number of significantly down- or up-regulated genes in head extracts of 27-day-old RU486-treated elavGS;HTTex1Q97 flies in comparison to head extracts of elavGS;HTTex1Q17 flies (|LFC| ≥ 1, p.adj < 0.01). **(c)** Venn-diagram displaying the numbers of significantly altered genes (|LFC| ≥ 1; p.adj < 0.01) and their overlap for all four treatment conditions. In red, the numbers of genes identified in at least two treatment conditions but not in the control condition (OFF) are highlighted. **(d, e)** String analysis of dysregulated protein clusters. Transcriptionally up- and downregulated protein clusters are exemplarily shown. See also the protein interaction network depicted in **Supplementary** Fig. 4. **(f)** Gene ontology (GO) enrichment analysis of DEGs identified under different experimental conditions (3d-ON/OFF, 6d-ON/OFF, 24d-ON) in RU486-treated HD transgenic flies compared to non-induced controls (p.adj < 0.05). **(g)** Cross-OMICS correlation analysis of proteins enriched in HTTex1Q97 aggregates and their associated downregulated mRNAs in head extracts of 24-day-old ON flies. Co-IP MS and RNAseq data obtained from elavGS;HTTex1Q97 and elavGS;HTTex1Q17 flies were compared (p.adj < 0.05). Overlapping genes encoding proteins with synaptic functions clustered with STRING (granularity score of 2) are shown. The fill color of each node corresponds to the log2FC from the IP-MS data, while the color of the node border represents the log2FC from the RNAseq data.

In order to define differentially expressed genes (DEGs), the transcriptome data sets (**Supplementary Table 3a**) of both genotypes (elavGS;HTTex1Q97 and elavGS;HTTex1Q17) were compared for all four experimental conditions (OFF, 3d-ON/OFF, 6d-ON/OFF and ON). In total, we identified 925 DEGs, of which 707 were abnormally up- and 225 down-regulated (|LFC| ≥ 1 and p.adj < 0.01) in heads of elavGS;HTTex1Q97 flies in comparison to elavGS;HTTex1Q17 controls (**Fig. 4b** and **Supplementary Table 3b**). We assumed that transcriptional changes detected under different experimental conditions may be more disease relevant than changes detected only once. For further network analysis, we therefore focused on expression changes (|LFC| ≥1 and p.adj < 0.01) that were detected in HD fly heads in at least two of the three experimental conditions (e.g., 3d-ON/OFF and 6d-ON/OFF; **Fig. 4c**). Through this approach, we defined 241 dysregulated genes, of which 171 were up- and 71 were downregulated (**Supplementary Table 3c**). Interaction network clustering with the STRING database ^54^ revealed 9 potentially dysregulated protein clusters with 5 or more nodes (**Supplementary** Fig. 4), indicating that progressive mutant HTTex1Q97 aggregation in neurons is associated with transcriptional dysregulation of a large number of subcellular processes. This includes a transcriptional up-regulation of processes such as DNA replication (**Fig. 4d**) or the transcriptional downregulation of key synaptic processes such as the release of neurotransmitters (**Fig. 4e**). Very similar results were obtained when DEGs (p.adj < 0.05) without a predefined LFC cut-off (**Supplementary Table 3d**) were investigated. An analysis of enriched GO terms confirmed that processes involved in neurotransmission and trans-synaptic signalling were transcriptionally downregulated in brains of HD transgenic flies with neuronal HTTex1Q97 aggregates in comparison to non-HD controls (**Fig. 4f and Supplementary Table 3e**). By comparing OMICS data sets, we defined a set of 93 proteins that are co-enriched with HTTex1Q97 aggregates (**Fig. 3c and 3d**) and whose transcripts levels are significantly decreased in heads of elavGS;HTTex1Q97 flies with amyloid-like mHTTex1 aggregates (24d-ON; **Fig. 4a**). Utilizing STRING followed by a gene set enrichment analysis, we defined transcriptionally dysregulated protein clusters with potential functional relevance in neurons. This analysis revealed a highly connected cluster of key synaptic proteins such as Vmat, Syn (synuclein) or nSyb (synaptobrevin) that are significantly enriched in HTTex1Q97 aggregates while their transcripts are downregulated (**Fig. 4g**). These studies support our hypothesis that the sequestration of key synaptic proteins into mHTTex1 aggregates (**Fig. 3d-3h**) and their transcriptional downregulation (**Fig. 4f**) may cause synaptic dysfunction and neurotoxicity in HD transgenic flies.

### Machine learning-based analysis of transcriptome and lifespan data reveals genes encoding synaptic proteins as positive predictors of fly survival

We hypothesized that at least a fraction of the HTTex1Q97 aggregation-induced transcriptome changes detected with RNA-seq in fly head extracts might be predictive of survival (**Fig. 4b**). Next, different machine learning-based computational models were trained with available gene expression (**Supplementary Table 4**) and survival data (**Supplementary Table 5)** to predict the lifespans of HD (elavGS;HTTex1Q97) and control (elavGS; HTTex1Q17) fly strains raised under different experimental conditions on food with or without the hormonal inducer RU486 (**Fig. 4a**). Cross-validation was then applied to compare the performance of two machine learning methods: elastic net (EN) regression and random forest (RF) regression (**Fig. 5a**). We found that EN regression models outperform RF regression models, when different evaluation measures such as RMSE (root mean squared error), MAE (mean absolute error), cosine similarity or Kendall’s coefficient were considered (**Supplementary** Fig. 5a). Based on these measures, EN regression models were finally applied for the selection of “lifespan-predicting genes” from gene expression data sets. Here, we focused on genes that were overlapping between all eight EN models from the cross-validation and were reliably identified as predictors irrespective of whether all genes or only subsets of top varying genes were given as input features (**Supplementary** Fig. 5b). Using this approach, we robustly identified 9 genes that are positively related to lifespan (i.e., genes whose expression is high when flies have a long lifespan **Fig. 5b**). In addition, two negative lifespan-predictive genes were identified (i.e., genes whose expression is high in flies with a short lifespan; **Fig. 5c**). For reasons of practicability, we next focused our efforts on the characterization of positive lifespan-predictive genes. The putative influence of negative lifespan predictors on fly survival will be assessed in an independent follow-up study.

**Figure 5:**
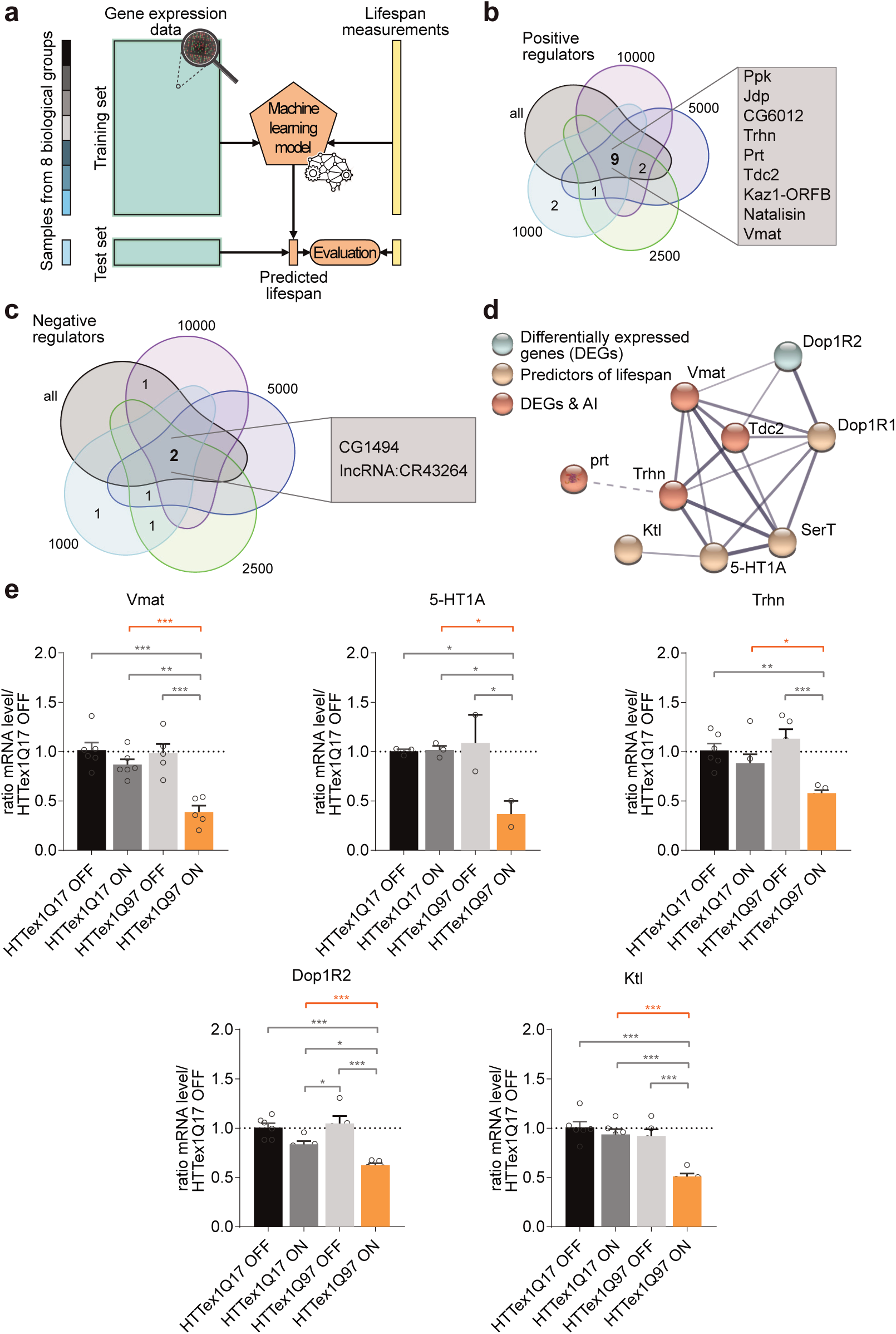
Machine learning-based analysis of transcriptome and *Drosophila* lifespan data reveals positive predictors of fly survival. **(a)** Workflow of the machine learning-based data analysis strategy. Eight different biological groups defined by treatment conditions (duration of transgene induction with RU486) and genotypes (elavGS;HTTex1Q17 or elavGS;HTTex1Q97) were defined. By utilizing a cross-validation approach and both gene expression and lifespan measurements as input data lifespan predictive genes were defined. **(b, c)** Venn diagrams showing positive (B) and negative (C) *Drosophila* lifespan-predicting genes and their overlap when EN regression was performed with all genes or different subsets of the top k varying genes (k=1000, 2500, 5000, 10000). The predicted positive and negative lifespan-predicting genes are displayed (grey box). **(d)** Expanded network of genes involved in monoamine signaling and/or biosynthesis. The blue node was identified by DEG analysis; brown nodes were obtained by machine learning-based data analysis. Nodes marked in red indicate a core cluster; they were identified by both computational data analysis strategies (DEG-based protein interaction clustering and EN regression). **(e)** Quantification of transcript levels of 5 selected genes encoding proteins involved in neurotransmitter signalling validates reduction of mRNAs. qPCR was performed with head extracts of 27-day-old HD transgenic flies after initiation of transgene expression with RU486 in 3-day-old flies (24d ON, also see Fig. 4a). The actin gene was used as a reference gene for qPCR analysis. Results of independent experiments are depicted as white dots and as mean ± SEM of biological replicates; statistical significance was assessed by One-way ANOVA Tukey‘s post-hoc test.

Among the selected positive lifespan predictors (**Fig. 5b**), four genes (*Trhn*, *prt*, *Tdc2* and *Vmat*) encode proteins that control the synthesis, transport and release of neurotransmitters in fly neurons ^55,56^, suggesting that efficient communication at synapses is critical for fly survival. For example, the genes *Trhn* and *Tdc2* encode enzymes (tryptophan hydroxylase and tyrosine decarboxylase 2) that play key roles in the biosynthesis pathways of serotonin and octopamine, respectively ^57,58^. Neuronal production of these neurotransmitters is of critical importance for locomotor activity and reproductive development of flies ^55^. In comparison, *Vmat* encodes a well-conserved vesicular monoamine transporter (Vmat) that facilitates the uptake of neurotransmitters such as serotonin and dopamine into synaptic vesicles. In synapses, this process is critical for the efficient release of multiple neurotransmitters into the synaptic cleft ^59^. A similar function was also suggested for the gene *prt*, which encodes the neuronal protein portabella (Prt). Similar to Vmat, this protein likely functions as a vesicular transporter in *Drosophila* neurons ^56^. Strikingly, an association of the proteins Vmat, Tdc2, and Trhn was observed when dysregulated fly transcripts were analyzed with the STRING tool (**Fig. 5d**), confirming that their cellular functions are linked. Together, these investigations suggest that synaptic processes that control the production and release of neurotransmitters in neurons (**Fig. 4e and 4f**) are critical for the survival of HD flies.

Finally, we applied an established qPCR method to quantify the transcript levels of 5 selected genes (*5-HT1A*, *Dop1R2*, *Ktl*, *Trhn*, *Vmat*), which all play a role in neurotransmitter biosynthesis and/or signaling ^60^ and whose expression in HTTex1Q97 RNAseq data sets was significantly decreased compared to HTTex1Q17 data sets (**Supplementary Table 3d**). Transcript levels of the selected genes were quantified in fly head extracts prepared of 27-day-old elavGS;HTTex1Q97 and elavGS;HTTex1Q17 flies, which were propagated for 24 days on food with or without the inducer RU486 (**Fig. 4a**). We measured significantly lower *Vmat, 5-HT1A*, *Trhn, Dop1R2* and *Ktl* transcript levels in head lysates of RU486-treated elavGS;HTTex1Q97 flies in comparison to RU486-untreated elavGS;HTTex1Q97 flies (**Fig. 5e**), supporting the hypothesis that formation of amyloid-like HTTex1Q97 aggregates in fly neurons is critical for the transcriptional downregulation of key synaptic genes.

### Knockdown of *Vmat* transcripts with RNAi decreases the survival of transgenic HD flies with neuronal amyloid-like HTTex1Q97 aggregates

Previous studies indicate that Vmat facilitates the uptake of neurotransmitters into vesicles in serotonergic, dopaminergic and octopaminergic neurons in *Drosophila* ^59^. We assumed that an siRNA-mediated downregulation of *Vmat* transcripts might further shorten the lifespan of elavGS;HTTex1Q97 HD flies that contain neurotoxic HTTex1Q97 aggregates, as this important synaptic function may be required for fly survival. To perform RNAi knockdown experiments, we generated the elavGS;HTTex1Q97;Vmat(KD) and elavGS;HTTex1Q17;Vmat(KD) fly strains that co-express HTTex1Q97 or HTTex1Q17 together with a *Vmat* mRNA-targeting RNAi molecule ^61^ in neurons through the Elav-GAL4 driver. We first quantified *Vmat* transcript levels in head extracts of 6-day-old RU486-treated HD flies by qPCR. As shown in **Fig. 6a** and **Supplementary** Fig. 6a, we measured significantly lower *Vmat* transcript levels in heads of RU486-treated elavGS;HTTex1Q97;Vmat(KD) and elavGS;HTTex1Q17;Vmat(KD) fly strains that express *Vmat* transcript-targeting siRNAs than in non-siRNA-expressing control flies (elavGS;HTTex1Q97 and elavGS;HTTex1Q17), confirming the knockdown of *Vmat* gene expression. Then, we quantified the survival of RNAi and non-RNAi-expressing HD flies. Strikingly, we determined a median lifespan of ∼29 and ∼23 days for RU486-treated elavGS;HTTex1Q97 and elavGS;HTTex1Q97;Vmat(KD) flies, respectively (**Fig. 6b and 6c**). This indicates that knockdown of *Vmat* expression with siRNA significantly decreases the lifespan of HD transgenic flies that progressively accumulate amyloid-like HTTex1Q97 aggregates in neurons (**Fig. 3b**). Interestingly, the median lifespan of RU486-treated elavGS;HTTex1Q17;Vmat(KD) flies was not significantly shortened in comparison to elavGS;HTTex1Q17 control flies upon neuronal siRNA expression (**Supplementary** Fig. 6b and 6c), indicating that a knockdown of *Vmat* transcript levels with siRNA molecules in healthy neurons is insufficient to significantly reduce survival.

**Figure 6:**
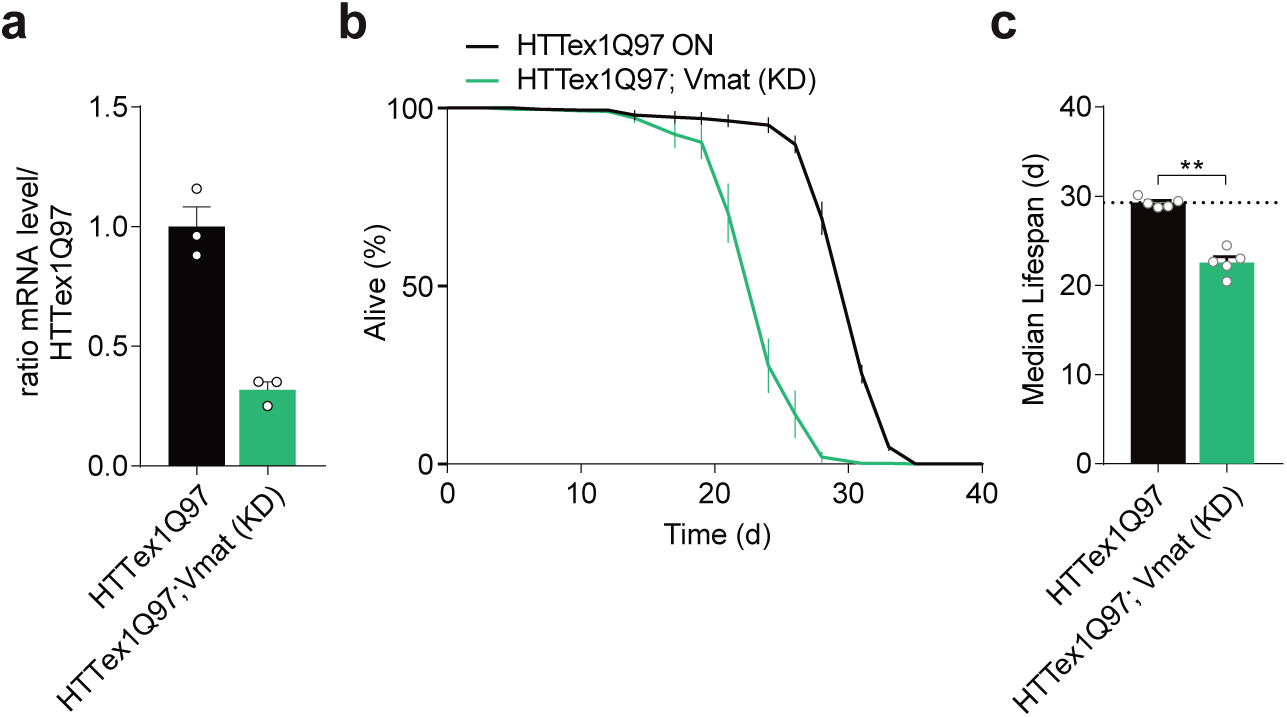
Knockdown of Vmat with RNAi decreases the survival of HD transgenic flies with neuronal amyloid-like HTTex1Q97 aggregates. **(a)** Quantification of *Vmat* transcript levels in brains of non-RNAi- and RNAi-expressing elavGS-driven HD (Q97) flies. Transgene expression was induced with RU486 in 3-day-old flies and samples were analysed after 6 days. Results of three biological replicates are depicted as white dots and as mean ± SEM; statistical significance was assessed by One-way ANOVA Tukey‘s post-hoc test. **(b)** Lifespan analysis of non-RNAi and RNAi-expressing elavGS;HTTex1Q97 flies after induction of transgene expression with RU486 in 3-day-old animals. Lifespan is plotted as the percentage of surviving flies of 5 biological replicates (∼50-100 flies each). Results are depicted individually as mean ± SEM of biological replicates. **(c)** Median life span calculated from survival curves in (**b)**. Average survival of each experiment (n ≥ ∼50 flies) is presented as white dots. Bars are mean ± SEM from 5 independent experiments; statistical significance was assessed by One-way ANOVA Dunnett’s post-hoc test; data were compared to RU486-treated elavGS;HTTex1Q97 transgenic flies.

## DISCUSSION

The accumulation of short N-terminal HTT fragments with pathogenic polyQ tracts in neuronal inclusion bodies (IBs) is a pathological hallmark of HD patient brains ^1,62^. Whether the aggregation of mHTT fragments in IBs is a neuroprotective process or rather contributes to the development of disease symptoms in HD patients and transgenic mouse models ^63^, however, is not well understood. In this study, we investigated the relationship between mHTTex1 aggregation in *Drosophila* neurons, the sequestration of proteins into mHTTex1 aggregates, transcriptional dysregulation and mortality of HD transgenic flies, to assess the impact of mHTTex1 aggregate species on disease development. We focused our efforts on pathogenic variants of mHTTex1, which *per se* have a high aggregation propensity ^6,64^ and are produced by abnormal splicing both in patients and in a HD knock-in mouse model ^18^. Thus, mHTTex1 fragments are likely to play a pathogenic role in HD, independently of full-length mHTT and N-terminal fragments released by proteolytic cleavage from the full-length protein ^18^. In this study, we found that a mHTTex1 fragment with a very long pathogenic polyQ tract (HTTex1Q97) rapidly forms SDS-stable, amyloid-like protein aggregates in neurons of very young HD flies (**Fig. 1d**), while the aggregation of a similar fragment with a shorter pathogenic polyQ tract (HTTex1Q49) was much slower and only detectable much later in older animals. Strikingly, HD flies, in which the amyloidogenic HTTex1Q97 aggregates appeared early in neurons, died significantly earlier than the HD flies, which accumulated HTTex1Q49 aggregates later in life (**Fig. 1e and 1f**). This indicates that mHTTex1 aggregation in neurons and fly mortality are causally linked. The rate of mHTTex1 aggregation, which is determined by the length of the pathogenic polyQ tract ^47^, dictates premature fly death. Independent studies with additional mHTTex1 protein variants support these conclusions. Comparing a mHTTex1 protein with a long uninterrupted polyQ tract (Ex1Q75), which rapidly forms amyloid-like protein aggregates in fly neurons (**Fig. 2b**), with mHTTex1 variants Ex1Q61ENK14 and Ex1Q71P4 with polyQ interruptions and reduced aggregation propensities, we found the latter to be significantly less neurotoxic to HD transgenic flies (**Fig. 2d and 2e**).

Using an RU486-inducible expression system ^38^ in young adult flies, we could show that amyloid-like HTTex1Q97 aggregates progressively accumulate over a relatively long time in fly neurons (∼20 days) without significantly affecting fly survival (**Fig. 3b**). This indicates that the immediate neurotoxicity of mHTTex1 aggregates is relatively low and that HD flies can cope with pathogenic amyloidogenic HTTex1Q97 aggregates for quite a long time (∼3-4 weeks) until they finally die rapidly within 3-5 days (**Fig. 3b**). This indicates that amyloidogenic mHTTex1 aggregates need to persist in neurons for some time until sufficient neurotoxicity is caused and finally makes the biological system collapse. Our results are not altogether incompatible with previous observations in cell models indicating that the deposition of insoluble mHTTex1 aggregates in IBs is neuroprotective ^65^. However, neuroprotection only lasts for a limited time and can by no means prevent the premature death of HD transgenic flies (**Fig. 3b**). We propose that small, diffusible mHTTex1 fibrils accumulate in neurons of HD flies over time in addition to large IBs with HTTex1Q97 aggregates. These smaller structures have a much greater surface area in comparison to IBs that concentrate amyloidogenic mHTTex1 structures in a specific subcellular region ^66,67^ and thereby might prevent the interaction of amyloid-like mHTTex1 assemblies with other cellular proteins. Small, diffusible mHTTex1 fibrils might be more neurotoxic than IBs because they are more agile in spreading from cell-to-cell, leading to pathology in different brain areas ^68^. Once they are present in neurons at a critical concentration, they might interact abnormally with large numbers of proteins and thereby alter a multitude of critical neuronal functions at the same time, triggering a breakdown of proteostasis. Small fibrils have in fact been detected in the cytoplasm of neuronal cells with high-resolution imaging methods besides large IBs that contain insoluble mHTTex1 aggregates ^69,70^. Small, seeding-competent mHTTex1 fibrils were detected in brain extracts of HD mice after large IBs with insoluble aggregates were removed by high-speed centrifugation ^22,23^, supporting the hypothesis that such structures may play a pathogenic role in disease.

To assess the interactions of amyloid-like mHTTex1 aggregates with other cellular proteins, antibody-enriched co-immunoprecipitates were analyzed by MS. These studies revealed that synaptic proteins normally associated with vesicles or the active zone were co-enriched with pathogenic HTTex1Q97 aggregates, suggesting that the progressive accumulation of mHTTex1 aggregates perturbs critical synaptic functions in neurons. Co-localization studies in 3^rd^ instar larvae also indicated an association of HTTex1Q97 aggregates with synaptic proteins at NMJs (**Fig. 3f-3h**), supporting our proteomics and biochemical results in adult brains. We propose the following chain of events: In a first phase, fibrillar HTTex1Q97 structures accumulate at synapses through abnormal polyQ-mediated protein-protein interactions of mHTTex1 monomers ^41^. This leads to a sequestration of synaptic vesicle-associated proteins into amyloid-like mHTTex1 aggregates at synapses and a gradual perturbation of functional liquid protein condensates that are present at the presynaptic nerve terminals ^71^. In this phase, neurotoxicity of amyloid-like mHTTex1 aggregates in neurons is still relatively low. In a next phase, neurons react to this local accumulation of synaptic proteins at pathogenic mHTTex1 aggregates with a transcriptional response leading to a decrease in the expression of synaptic genes and finally to a severe impairment of synaptic transmission and premature death. Our observations that the expression of a large number of synaptic genes is reduced in brains of 27-day-old HD flies with neuronal HTTex1Q97 aggregates but not in controls are compatible with such a model (**Fig. 4f**). Transcripts encoding synaptic proteins were also shown to be significantly decreased in postmortem brains of HD patients ^72^, supporting the hypothesis that critical synaptic functions are impaired in HD brains. Previous evidence that pathogenic mHTTex1 fibrils can directly bind to lipid vesicles ^73^ and interact with different synaptic proteins in yeast two-hybrid and pull-down assays also supports our model ^74,75^. Cellular studies have previously shown that membranes of the endoplasmic reticulum or other subcellular organelles are indeed present in IBs with insoluble fibrillar mHTTex1 aggregates ^66,67^. Thus, it seems reasonable to speculate that the progressive formation of amyloid-like HTTex1Q97 aggregates in neurons over time causes a recruitment of membrane-associated synaptic proteins and concomitantly a decrease of their transcript levels, eventually leading to a significant alteration of neuronal communication, motor impairment and premature death of HD flies.

Through the application of machine learning methods for the analysis of available transcriptome and survival data sets, we were able to select “positive lifespan-predictive” genes, which are genes whose expression is high in neurons of flies with a long lifetime (**Fig. 5a and 5b**). Strikingly, this analysis revealed 3 genes (*Trhn*, *Tdc-2* and *Vmat)* that are closely linked functionally (**Fig. 5d**) and that together play a key role in *Drosophila* neurotransmission ^55^, supporting the hypothesis that synaptic functions are critical molecular determinants for the survival of flies. Importantly, our investigations with qPCR confirmed that *Trhn*, *Tdc-2* and *Vmat* transcript levels are indeed significantly decreased in heads of HD elavGS;HTTex1Q97 flies that contain large amounts of amyloid-like HTTex1Q97 aggregates and that die prematurely after ∼32 days. In comparison, control flies without neuronal HTTex1Q97 aggregates have a median lifespan ∼80-90 days. Knocking down *Vmat* expression with siRNA in neurons even further shortened the lifespan of elavGS;HTTex1Q97 flies in comparison to non-siRNA-expressing elavGS;HTTex1Q97 control flies (**Fig. 6a-6c**), substantiating the hypothesis that the synthesis and release of neurotransmitters such as dopamine and serotonin are critical for the survival of HD flies. A neuroprotective function of the related protein VMAT2 in human brains has previously been reported ^76^ and reduced levels of monoaminergic neurotransmitters such as dopamine were indeed measured in postmortem brains of HD patients ^77^. Earlier investigations also demonstrated that dopamine and serotonin levels are significantly reduced in striatal tissues of R6/2 HD transgenic mice ^78^, supporting the relevance of our observations in transgenic HD flies.

Our *in vivo* studies with HD transgenic flies have important implications for a better understanding of the disease process and the development of potential novel disease-modifying therapies for HD. First, our investigations support the hypothesis that formation of amyloid-like, self-propagating mHTTex1 aggregates in neurons is a pathogenic process that causes neurotoxicity, motor impairment and premature death of HD flies over time. More in-depth research and development efforts should be undertaken to identify mHTTex1 aggregation modulating small molecules and to assess their effects on aggregation and neurotoxicity in other suitable *in vivo* model systems such as HD mice. We propose that therapeutic molecules that efficiently slow down mHTTex1 polymerization or promote mHTTex1 degradation should also reduce neurotoxicity and significantly extend the lifespan of HD transgenic flies. Previous reports suggested that HD is a synaptopathy ^79^, meaning that the impairment of critical synaptic processes or a loss of synapses plays an important role in the development of disease symptoms. Our studies support such a disease mechanism for HD. Our data also suggest that treatment of HD patients with tetrabenazine, a well-characterized VMAT2 inhibitor ^80^ generally applied to reduce motor symptoms, may cause neurotoxicity in late-stage, akinetic HD patients, when dopamine levels are already very low^77^. Therapeutic molecules that increase VMAT2 expression or activate VMAT2 activity in neurons may improve disease symptoms and extend the lifespan of HD patients. Further, detailed investigations with compounds that increase neurotransmission or reverse the transcriptional downregulation of synaptic proteins in HD models will be necessary to address this question.

## METHODS

### EXPERIMENTAL MODELS

#### Fly generation and husbandry

*Drosophila melanogaster* strains were reared under standard laboratory conditions and raised at 25°C and 70% humidity on semi-defined medium (Bloomington recipe). For HTTex1 expression experiments with Elav-GAL4/X or elavGS-GAL4 (GS)/X flies were kept at 25 or 29°C. For all experiments with adult flies animals were propagated at 25°C. Experiments with larvae were performed at 29°C. Vmat siRNA flies (BloomStockNo. 44471) were obtained from the Bloomington *Drosophila* Stock Center. The w1118 lines was provided by R.P. Zinzen (MDC, Berlin) or by the Bloomington *Drosophila* Stock Center. The generation of HTTex1Q17- or HTTex1Q97-expressing HD flies was previously described ^23^. Transgenic flies were generated through cloning of cDNAs encoding HTTex1Q49, HTTex1Q0, Ex1Q17, Ex1Q75, Ex1Q61ENK14 and Ex1Q71P4 into pUAST-attB-rfA and subsequent site-directed insertion into the third chromosome (68E) using the PhiC31 integrase at Rainbow Transgenic Flies Inc. (Camarillo, CA, USA). Transgenic lines were crossed with the balancer strain (CyO/Sp;TM6,Tb/MKRS,Sb) to produce stable fly lines. All *Drosophila* strains were cultured on standard medium at 25°C and 65% humidity with a 12 h light-dark cycle. ElavGS-GAL4-mediated transgene expression was induced by culturing flies on standard medium with 400 µM RU486 (Mifepristone).

### IMMUNOSTAININGS OF LARVAE

#### Immunohistochemistry for confocal and STED microscopy

Immunohistochemistry dissections were performed in haemolymph-like solution 3 (HL3); composition in mM: 70 NaCl, 5 KCl, 20 MgCl_2_, 10 NaHCO_3_, 5 trehalose, 115 sucrose, 5 HEPES, pH adjusted to 7.2) by opening the 3^rd^ instar larvae dorsally along the midline and removing the entrails. Dissections were fixated with 4 % paraformaldehyde in PBS (pH 7.2) for neuromuscular junction (NMJ) staining for 10 min. After fixation, the filets were washed with PBS plus 0.05 % Triton-X 100 (PBT) and blocked for 60 min in 5 % normal goat serum (NGS, Sigma Aldrich S2007). For immunostainings, the larvae were incubated with primary antibodies at 4° C overnight and subsequently washed in a 0.05 % PBT solution for 2 h at room temperature (RT). Larvae were then incubated for 2-3h with secondary antibodies at RT. Washing procedures were repeated. Larvae were finally mounted in Mowiol (Sigma Aldrich). Primary and secondary antibodies used in these studies are shown in key resources table.

#### Image acquisition and analysis

Conventional confocal images were acquired with Leica DMI 6000 (SP8) (Leica Microsystems). For confocal scans, a HC PL APO CS2 63x /1.40-N.A. oil objective (Leica Microsystems) was used. Images were acquired at approximately 20°C and fluorochromes are indicated in the antibody section described above. Imaging medium was immersion oil (Sigma-Aldrich 55822). For signal detection HYD (high-sensitive) 400-800 nm spectral descanned for green and red channels and PMT 400-800 nm spectral descanned for far red channels were used for confocal scans. The NMJ z-stacks had a step size of 0.2 – 0.3 μm between single optical slices. All images were acquired using the LAS X software (Leica Microsystems, Wetzlar, Germany). The ImageJ 1.52n software was used for the analyses of confocal images. GraphPad PRISM, version 5.01 was used for statistical analyses. Images for figures were processed, if necessary, with ImageJ software to enhance brightness using the brightness/contrast function. Confocal stacks were processed with the image software Fiji (http://fiji.sc) ^81^.

#### NMJ quantifications

Confocal stacks were analyzed with the image software Fiji (http://fiji.sc) ^81^. Image analysis followed the standard protocol as described by ^82^. Type1b neuromuscular junctions (NMJs) on muscle 4 were analyzed. The original confocal z-stacks were converted to maximal projections. The HRP-Cy5 antibody signal was used as the template for a mask, restricting the quantified area to the shape of the NMJ. Within the HRP mask the mean pixel intensity within the NMJ area measured for the different channels. N represents 1-2 NMJs from 5-6 different animals/genotype.

#### Line graphs of confocal images

After the creation of a maximum projection a line was placed through the chosen and representative bouton, saved as ROI. The ROI manager MultiPlot tool was used to determine peak to peak distance and plotted within GraphPad Prism v8.01.

#### Quantification and statistical analysis

Data were analyzed using GraphPad Prism v8.01. Normality was tested with the D’Agostino & Pearson omnibus normality test. If data were normal distributed two groups were compared, an unpaired, two-tailed t-test was used (**Fig. 3f-3h and Supplementary** Fig. 3f**-3h**). *p<0.05, **p<0.01, ***p<0.001.

### METHOD DETAILS

#### Cloning of expression vectors

To generate the plasmids encoding GST-Ex1Q23 and GST-Ex1Q48 the phosphorylated oligonucleotides HTT123_A and _B were annealed to generate the coding sequence for the V5-tag with single stranded 5’ and 3’ overhangs for XhoI and NotI cleavage sites. Linearization of the plasmids pGEX-6P1-Ex1Q23 and pGEX-6P1-Ex1Q48 using XhoI and NotI and ligation of linearized plasmid DNA to the annealed oligonucleotides generated the plasmids pGEX-6P1-Ex1Q23 and pGEX-6P1-Ex1Q48. In order to generate expression plasmids for the V5-tagged protein variants of Ex1Q48, cDNAs encoding Ex1Q48 variants Ex1Q38ENK10 and Ex1Q45P3 with the respective amino acid exchanges were produced by gene synthesis (GeneArt® Gene Synthesis service, Thermo Fisher). cDNAs were PCR-amplified with the primers HTT123_C and _D. The primer pair was designed to introduce a DNA sequence encoding a C-terminal V5-tag and additional endonuclease cleavage sites upstream (EcoRI) and downstream (NotI) of the coding sequence. The resulting PCR products were digested using the EcoRI and NotI endonucleases and cloned into the plasmid pGEX-6P1-Ex1Q48 after excision of the fragment encoding the Ex1Q48 proteins with EcoRI and NotI.

#### Cloning of fly vectors

To generate HD transgenic flies cDNAs encoding the proteins HTTex1Q0, HTTex1Q17, HTTex1Q49 or HTTex1Q97 were transferred from the gateway-compatible pDONR221 vectors into the destination vector pUAST-attB-rfA ^39^, resulting in the creation of the plasmids pUAST-HTTex1Q0, -HTTex1Q17, -HTTex1Q49 or -HTTex1Q97. In order to express variants of mutant HTTex1 in flies, cDNA fragments encoding Ex1Q75, Ex1Q61ENK14, Ex1Q71P4 or Ex1Q17 were produced by gene synthesis (GeneArt® Gene Synthesis service, Thermo Fisher). cDNAs were PCR-amplified with the primers HTT123_G and _H. The primer pair was designed to introduce DNA sequences encoding a C-terminal V5-tag and attB recombination sites upstream (attB1) and downstream (attB2) of the coding sequence. To generate entry plasmids the resulting PCR products were integrated into pDONR221 (Thermo Fisher) through a BP recombination reaction. LR recombination reactions were performed to shuttle cDNAs into the destination vector pUAST-attB-rfA ^39^. The correct identities of all plasmids were confirmed by Sanger sequencing using the sequencing primers pUAST-fw and pUAST-rev.

#### Recombinant protein expression

The GST (N-terminal) and V5-tagged (C-terminal) HTTex1 fusion proteins (Ex1Q23, Ex1Q48, Ex1Q38ENK10 and Ex1Q45P3) were produced in *E. coli* BL21-CodonPlus-RP and affinity-purified on glutathione-sepharose beads. Purified proteins were dialyzed over night at 4 °C against 50 mM Tris-HCl pH 7.4, 150 mM NaCl, 1 mM EDTA and 5% glycerol, snap-frozen in liquid N_2_ and stored at -80 °C. Protein concentrations were determined with a NanoDrop spectrophotometer. Prior to use, protein solutions were ultra-centrifuged at 190,000 x g for 40 min to remove aggregated material.

#### *In vitro* protein aggregation

Spontaneous aggregation GST-HTTex1 fusion proteins with interrupted and non-interrupted polyQ tracts was initiated by addition of 14 U PreScission protease (GE Healthcare) per nmol purified GST fusion protein (4 µM). The aggregation reactions were performed in 50 mM Tris-HCl pH 7.4, 150 mM NaCl, 1 mM EDTA and 1 mM DTT at 25 °C and constant agitation (450 rpm) for up to 168 hours.

#### Filter retardation assays (FRAs)

FRAs were performed as described previously ^83^. Briefly, equal volumes of 500 ng of protein samples and 4 % SDS solution with 100 mM DTT were mixed and boiled at 95 °C for 5 min. By applying vacuum, samples were filtered through a cellulose acetate membrane with 0.2 µm pores (Schleicher and Schuell, Germany) and washed twice with 100 µl 0.1 % SDS. For analysis of tissue homogenates, 15 or 75 µg of total protein prepared from *Drosophila* heads were filtered per dot. Membranes were blocked with 5% skim milk in PBS/0.05 % Tween20 (PBS-T) for at least 30 min. Aggregates retained on the membrane were detected using an anti-HTT or an anti-V5 antibody followed by an appropriate peroxidase-coupled secondary antibody. Signals were quantified using the AIDA image analysis software (Raytest, Germany).

#### SDS-PAGE and immunoblotting

Samples of HTTex1 aggregation reactions with recombinant proteins were mixed with loading buffer (50 mM Tris-HCl pH 6.8, 2% SDS, 10% glycerol and 0.1% bromophenol blue) and boiled at 95 °C for 5 min. Samples were loaded onto Novex NuPAGE 4-12% Bis-Tris gradient gels (Life Technologies). For analysis, 10 µg of total protein prepared from *Drosophila* heads were loaded per lane. SDS-PAGE and immunoblotting were performed according to manufacturer recommendations. Immunodetection was performed with the antibodies indicated for the respective experiment. Signals were quantified using Fiji (ImageJ). Proteins separated in gels were blotted onto a nitrocellulose membrane (0.45 μm) in NuPAGE transfer buffer (2×, 10% methanol) using a Power Blotter, at 20 V constant voltage for 1 h. Then, the nitrocellulose membrane was shortly washed with PBS-T and stained with Ponceau S solution. Images of the stained nitrocellulose membranes were taken using the integrated Ponceau S channel in the iBright CL1500. The staining was removed by a short incubation with PBS-T (20 ml). Then, the membrane was blocked in 3% milk-PBS-T (20 ml) for 1 h at RT and incubated with antibodies (PHP2, V5, S830, MAB5492, Brp or Vmat) diluted in 3% skim milk-PBS-T (10 ml) overnight at 4°C. The next day immunoblots were washed three times with PBS-T (20 ml) for 10 min at RT. Corresponding mouse, rabbit or sheep HRP-conjugated secondary antibodies were diluted in 3% milk-PBS-T (10 ml) and applied to the immunoblots for 1 h at RT. Immunoblots were finally washed three times with PBS-T (20 ml) for 10 min at RT. For immunodetection of proteins blots were incubated with WesternBright Quantum chemiluminescence substrate (8 ml) for 2 mins and then imaged with the iBright CL1500 in the appropriate channels.

#### Immunoprecipitation

Dynabeads^TM^ protein G (37.5 μl) were washed two times with PBS and finally resuspended in PBS (350 μl). Then, the anti-HTT antibody PHP2 (12 μl) was added to the beads and the mixture was incubated for 30 min at RT on a rotating wheel. Finally, the supernatant was discarded and the beads were washed two times with PBS (350 μl). For immunoprecipitation fly head lysates were added to the beads, diluted with BLB buffer up to a total volume of 500 μl, and incubated overnight on a rotating wheel. The next day, the flow-through (FT) was discarded and the beads were washed with BLB (1× 2 min and 2× 1 min) and proteins eluted with NuPAGE LDS sample buffer (1×) and DTT (50 mM) through boiling at 95°C for 5 min. For subsequent LC-MS/MS applications, all volumes were doubled, and the last two washing steps were performed with 10 mM Tris-HCl pH 7.4 buffer (1.2 ml). Subsequently, beads were separated into two samples and one-half of the pelleted beads was snap-frozen in liquid N_2_ and finally submitted for LC-MS/MS measurements. The other half of the pelleted beads was analyzed by SDS-PAGE and immunoblotting.

#### Atomic force microscopy (AFM)

Purified GST-tagged Ex1Q23, Ex1Q48, Ex1Q38ENK10 or Ex1Q45P3 fusion proteins (4 µM) were incubated with Prescission protease (PSP) and spontaneous aggregation reactions were analyzed by AFM after 168 h. Aliquots of 15 µl were spotted onto freshly cleaved mica glued to a microscope slide. After incubation for 30 min to allow adsorption, samples were rinsed 4 times with 40 µl distilled water and dried over night at RT. Samples were imaged with a digital multimode Nanowizard II (JPK, Germany) atomic force microscope operating in intermittent-contact mode.

#### Image analysis of adult fly brains

Z projections of fly brain images were created using the maximum intensities of the separate stacks using ImageJ. Then, the red and green channels corresponding to the RBP ^51^ and HTT signals, respectively, were split. Using the JACoP plugin, intensity thresholds were manually set for the two channels and then the Manders’ coefficients (M1 and M2) were calculated ^84^. The Manders’ coefficients are based on the Pearson’s correlation coefficient and they provide a value between 0 and 1; with 0 representing non-overlapping pixels and 1 reflecting 100% overlap between the pixels from the two separate channels. For the analysis performed here, the M1 coefficient was used in order to reflect the summed intensities of green pixels for which the intensity in the red channel is over zero. The M1 coefficient therefore represents the amount of RBP signal overlapping with the HTTex1 signal over the total HTTex1 signal intensity. For the conditions of interest, the M1 coefficient was calculated for all fly brain images from the different biological replicates. After obtaining the M1 values, a bar graph visualizing the differences in the average M1 values was created, and statistical analysis using the ANOVA Kruskal-Wallis test was carried out on Graphpad Prism.

#### Genotyping of *Drosophila* strains

Genomic DNA was extracted using the DNeasy® Blood & Tissue Kit (Qiagen). Genomic DNA was PCR amplified with the genotyping primers HTTX and HTTY using the PWO DNA polymerase kit (Roche). PCR products were analyzed by agarose gel electrophoresis and DNA sequencing.

#### Reverse transcription and quantitative polymerase chain reaction (qPCR)

RNA was extracted from *Drosophila* heads using the RNeasy Kits (QIAGEN) and subsequent isopropanol precipitation. cDNA was synthesized using the High-Capacity cDNA Reverse Transcription Kit (Thermo Scientific) and qPCR was performed using the SYBR Green PCR Master Mix (Thermo Scientific). The primer pairs used for the targets HTTex1, 5-HT1A, Dop1R2, Ktl, Trhn, Vmat, actin and rp49 are shown in key resources table. SYBR Green analysis was performed using the ViiA7 Real-time PCR system (Thermo Scientific). The amount of mRNA detected was normalized to control actin or rp49 mRNA values.

#### Viability analysis of adult Drosophila melanogaster strains

Viability assays were performed through quantification of lethality of female transgenic flies. Flies were aged at 25°C and 65% humidity with a 12 h light-dark cycle, with 10 flies per vial. Flies were transferred onto new media every 3-4 days. Dead flies were recorded every 2-3 days. The number of flies per biological replicate are specified in the figure legend of the respective experiment. Median lifespan (age at which half of the tested population has died) was calculated by fitting survival curves to the log(inhibitor) vs. normalized response (variable slope) equation using GraphPad Prism.

#### Analysis of motor performance (climbing assay)

Ten female flies were placed in a closed empty vial and gently tapped to the bottom of the vial. The percentage of flies that climbed 8 cm within 15 sec was recorded. Flies were aged at 25°C and 65% humidity with a 12 h light-dark cycle (10 flies per vial) and were monitored and transferred twice a week. The number of flies per biological replicate are specified in the figure legend of the respective experiment.

#### Analysis of circadian rhythm and activity

Locomotor activity was monitored using the *Drosophila* Activity Monitoring (DAM) System as previously described ^85^. Male flies were placed individually into glass locomotor-monitoring tubes (65 mm x 5 mm) containing standard medium. The glass tubes were sealed using cotton buds. The tubes were inserted into the sensor system. Flies were aged at 25°C and 65% humidity with a 12 h light-dark cycle. Within the sensor system three infrared beams are directed through each glass tube. Activity of transgenic flies was assessed by recording the number of beam breaks within a 5 min interval. The number of flies per biological replicate are specified in the figure legend of the respective experiment.

#### Preparation of *Drosophila* head lysates for FRAs

*Drosophila* head lysates were produced by homogenizing fly heads in Tris-HCl pH 7.4, 0.8 M NaCl, 1 mM EDTA, 10% sucrose, 0.25 U/µl benzonase and complete protease inhibitor cocktail using the Precellys Evolution homogenizer (CK14 vials, 3 x 10 sec intervals of shaking at 6,000 rpm with 10 sec break). Lysates were centrifuged for 10 min at 8,000 rpm (4°C). The supernatant was transferred to a new tube and total protein concentration was determined with a Pierce™ BCA assay using BSA as a standard.

#### Dissection and immunostaining of *Drosophila* adult brain

Whole brains of adult flies were dissected in ice-cold haemolymph-like saline (HL3) solution ^86^, fixed for 20 min in 4 % paraformaldehyde (PFA) in PBS and permeabilized in PBS-T (1 % Triton^TM^ X-100) for 20 min at RT. Samples were blocked in 10 % normal goat serum (NGS) in PBS-T (0.3 % Triton^TM^ X-100) for at least two hours. Brains were incubated with the indicated primary antibody (1:500) in brain staining buffer (5 % NGS, 0.1 % NaN_3_ in PBS-T (0.3 % Triton^TM^ X-100) for 48 hours at 4 °C. Subsequently, brains were washed in PBS-T (0.3 % Triton^TM^ X-100) for 24 hours at 4°C with multiple buffer exchanges. Next, samples were incubated with appropriate secondary antibody in brain staining buffer for 24 h at 4 °C, washed six times for 30 min in PBS-T (0.3 % Triton^TM^ X-100) at RT and stored in VectaShield H-100 (Vector Laboratories) at least for one day at -20 °C. Brains were mounted and imaged using the Leica TCS SP8 confocal microscope. Images were analyzed using Fiji.

#### RNA sequencing and quantification of differential gene abundance

Total mRNA was extracted from *Drosophila* heads using TRIzol™ Reagent (Invitrogen). mRNA selection was performed by polyA enrichment and followed by cDNA library preparation using the Illumina TruSeq Stranded mRNA Library Prep Kit for high throughput (96 samples, 96 indexes). RNA-sequencing (RNA-Seq) was performed using Illumina NextSeq 500 by the Genomics core facility at the MDC (75 bp read length, paired-end sequencing and 150 cycles to gain a depth of 25 million reads/sample). RNA-seq reads were mapped to the *Drosophila* reference genome with STAR ^87^ version2.4.2a. Reads were assigned to genes using the FeatureCounts ^88^, Rsubread version v1.4.6-p5, using *Dm* BDGP6.92 gtf annotation. The differential expression analysis was carried out with DESeq2 ^89^, version 1.34.0. We used only protein-coding genes.

#### Label-free quantitative MS

Proteins enriched from frozen fly head extracts by co-IPs were analyzed at the Proteomics core facility at the MDC. Tryptic on-bead digestion was carried out following the protocol from Hubner et al ^90^ and then desalted on stage-tips ^91^. LC-MS measurements were carried out on an orbitrap Exploris 480 mass spectrometer coupled to an EASY-nLC 1200 system applying a 110 min gradient and operating the MS in data-dependent mode. For analysis, MaxQuant version 2.0.3.0 ^92^ was used employing MaxLFQ-based ^93^ quantitation. Carbamidomethylation was set as fixed and oxidized methionine as variable modification. For the Andromeda search, a Uniprot *Drosophila* database plus common contaminants was used in combination with the recombinant HTT sequences. Downstream analysis was done in R using two-sample moderated t-statistics with the limma package ^94^.

#### GO term enrichment analysis with RNA-seq data sets

Three sets of DEGs (3d-ON/OFF, 6d-ON/OFF, ON, **Supplementary Table 3D** were selected for a Gene Ontology (GO, RRID:SCR_002811, http://www.geneontology.org/) enrichment analysis (**Fig. 4f**), which was performed with the R package clusterProfiler ^95^. The enrichment of the shared GO term Biological Process in the DEG sets was computed as an exact p-value, FDR-corrected for multiple testing, of a given minimum hypergeometric (mHG) score determining the probability that the number of proteins in the DEG sets that map to the GO term is explained by chance alone. The gene ratios (**Fig. 4f**) are defined as n/N where n represents the number of genes of the given DEG set involved in the given biological process, while the divisor N indicates the number of genes of the given dataset associated with any GO term ^95^.

#### Identification of *Drosophila* lifespan-predicting genes

RNA-seq data were normalized as transcript per million (TPM) and log-transformed before further analysis (log2(TPM+1)). For combining RNA-seq data with lifespan data, we kept the individual RNA-seq replicates and assigned to each of them the lifespan mean value of the respective condition and genotype. To test the performance of different machine learning methods that predict lifespan from gene expression data, we performed leave-one-out cross-validation across the eight different biological groups defined by the combination of treatment condition and genotype. Four evaluation measures were used to compare predicted values and ground truth values. The root mean squared error (RMSE) is 0 for perfect predictions and greater than 0 otherwise. The cosine similarity is a length-normalized inner product of two vectors, yielding values between -1 and 1. Here, the first vector contains the pairwise differences of the predicted values to the training example values, and the second vector contains the pairwise differences of the ground truth values to the training example values. The cosine similarity is 1 if the predictions and ground truth values behave proportionally relative to the training examples, and -1 if they behave in the opposite way. Due to the limited size and variability of the test datasets, it is more informative to examine the predicted values in the context of the values from the training data than in isolation. Thus, we took difference vectors instead of prediction value vectors. Furthermore, we included two ranking-based measures. The mean absolute error (MAE) indicates the average absolute difference between the rank of the predicted value and the rank of the ground truth value within the combined set with the training values. Kendall’s rank correlation coefficient also uses the ranks of these combined sets to compute their correlation, ignoring tied values. We considered two types of machine learning approaches for lifespan prediction, random forests (RF) and elastic net regression (EN). The random forests were computed using the R package randomForest, version 4.7-1, setting the number of trees to 500 and the minimum node size to 3. The elastic net regression was done with the R package glmnet, version 4.1-3. The parameter alpha was set to 0.5. Input and output data were standardized with the mean and standard deviation of the respective training data. Predictions were taken at the central position of the default lambda sequence.

#### Bioinformatic data processing and visualization

Gene ontology (GO) terms of proteins were obtained using Metascape with the following parameters: p.adj < 0.01, overlap >3, enrichment >1.5 ^96^. Protein clusters of directly interacting proteins from protein lists were obtained by the Markov Cluster (MCL) algorithm ^97^ with a granularity score of 2 using the STRING app ^98^ within Cytoscape ^99^. Figures were created using Biorender.com (2023) in combination with Adobe Illustrator. For protein structure prediction the Alphafold 3 tool was applied. For investigating and visualizing of protein structures the PyMOL Molecular Graphics System was (Version 3.0 Schrödinger, LLC) was used.

### STATISTICAL ANALYSIS

Statistical parameters including the exact value of n, the definition of center, dispersion and precision measures (mean ± SEM or mean ± SD) as well as the statistical analysis chosen and statistical significance are reported in the figures and figure legends. Data is judged to be statistically significant when p <0.05 by the indicated statistical test. In figures, asterisks denote statistical significance as calculated by Student’s t test (*, p <0.05; **, p <0.01; ***, p <0.001). Statistical analysis was performed in GraphPad PRISM 7.

## Supporting information

Supplementary Figures, Supplementary Figure titles and legends and Table titles

## ACKNOWLEDGMENTS

This work was supported by the Hereditary Disease Foundation, USA (postdoctoral fellowship to A.A.); the CHDI Foundation, USA; iMed – Helmholtz Initiative on Personalized Medicine, Germany and the Helmholtz-Israel-Cooperation in Personalized Medicine, Germany (research grants to E.E.W.). We thank the Advanced Light Microscopy Technology Platform of Max-Delbrück-Center for Molecular Medicine, Berlin for general and technical support. We also thank the Genomics Technology Platform for conducting mRNA enrichment, library preparation and RNA sequencing. We wish to thank Dr. Alexander Buntru for his support and willingness to discuss biochemical questions. We are grateful to Prof. Stephan Sigrist for providing the destination vector pUAST-attB-rfA and to Prof. David Krantz for providing an anti-Vmat antibody. Finally, we thank Sigrid Schnoegl for discussing and revising the manuscript.

## AUTHOR CONTRIBUTIONS

A.A. and E.E.W. conceived the study and designed the experiments; A.A., E.G., C.H. L.R. and A.I. analyzed data. A.A., L.B. and S.B. performed in vitro experiments with protein variants of HTTex1. S.K. analyzed EM samples. A.A., L.B., F.S., L.R. and J.E. performed all studies with the HD transgenic flies. A.P. performed all studies with *Drosophila* larvae. M.B. and O.W.M.A performed immunostaining and confocal imaging of fly brains. L.R., O.P., and P.M. performed co-IP and MS experiments. C.L. performed the library generation and RNA-sequencing. A.A., E.G, J.E., A.I., M.P., C.H. and D.B. generated and analyzed RNA-sequencing data. A.A. and E.E.W. conceived and wrote the manuscript.

## MATERIALS AND CORRESPONDENCE

For the materials used, please see **Supplementary Table 6**.

CONTACT FOR REAGENT AND RESOURCE SHARING Further information and requests for reagents may be directed to and will be fulfilled by the corresponding author Erich E. Wanker.

