## Supplementary Figures, Supplementary Figure titles and legends and Table titles for "Formation of amyloid-like HTTex1 aggregates in neurons, downregulation of synaptic proteins and early mortality of Huntington’s disease flies are causally linked"

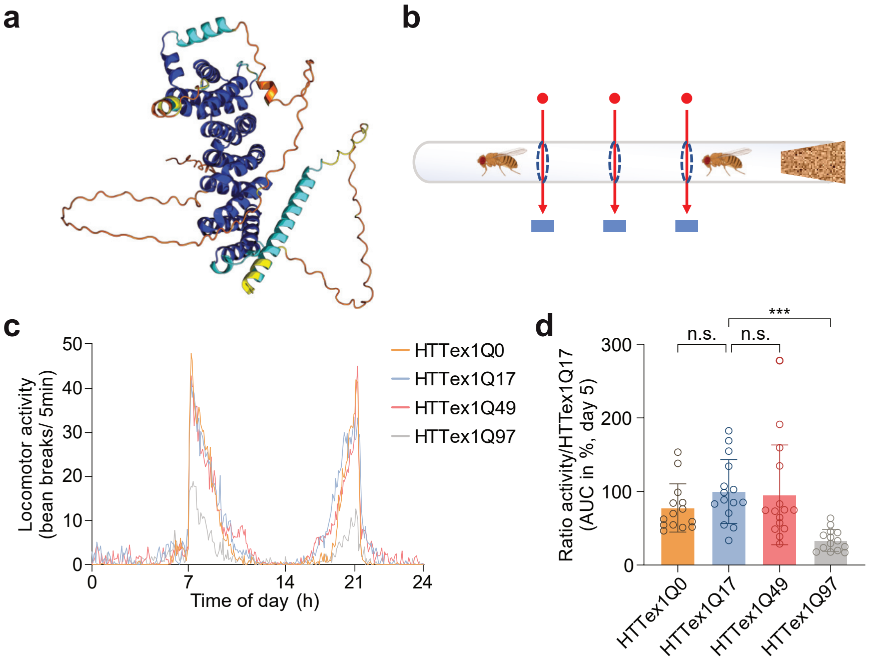


**Supplementary Figure 1**

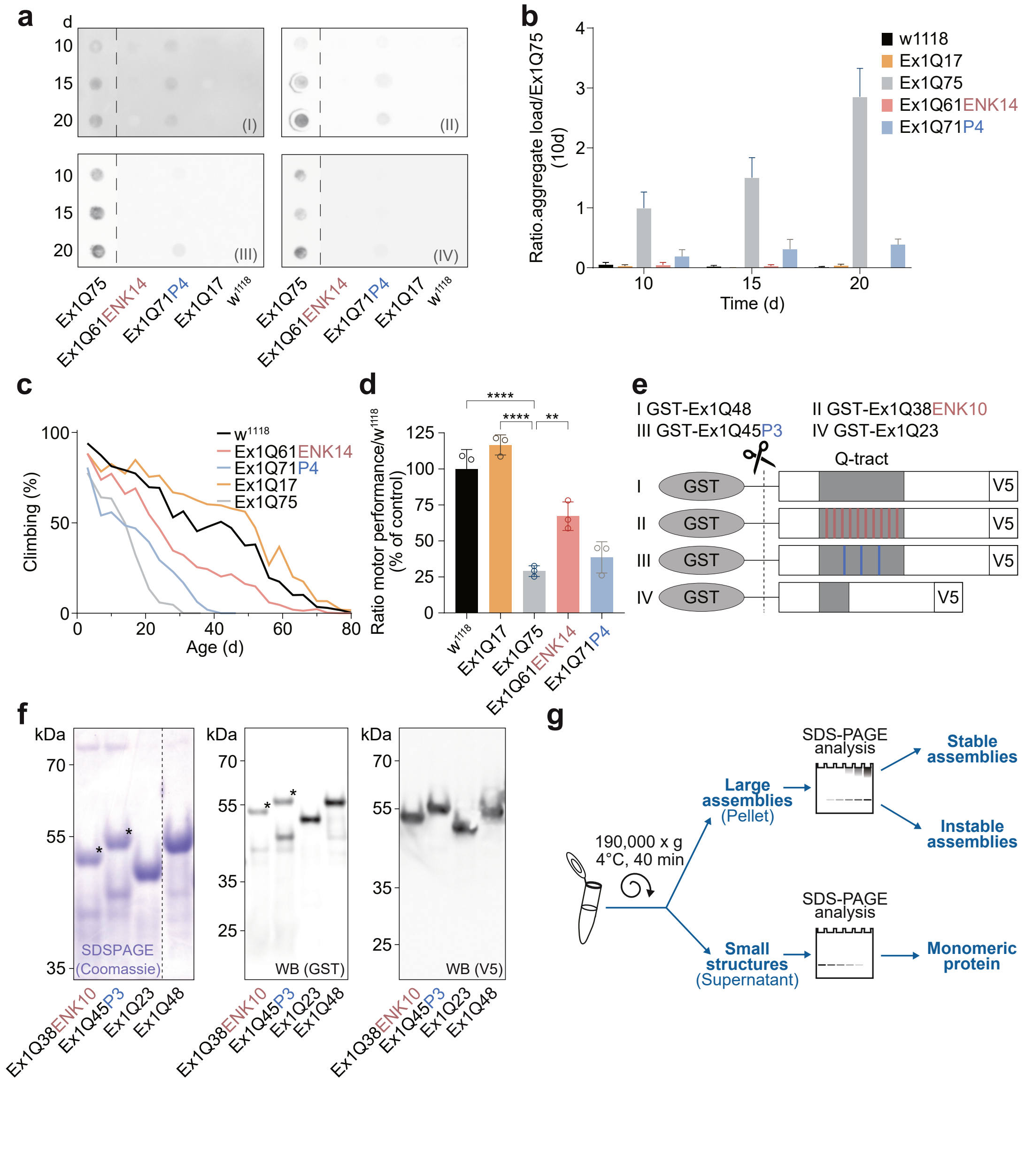


**Supplementary Figure 2**

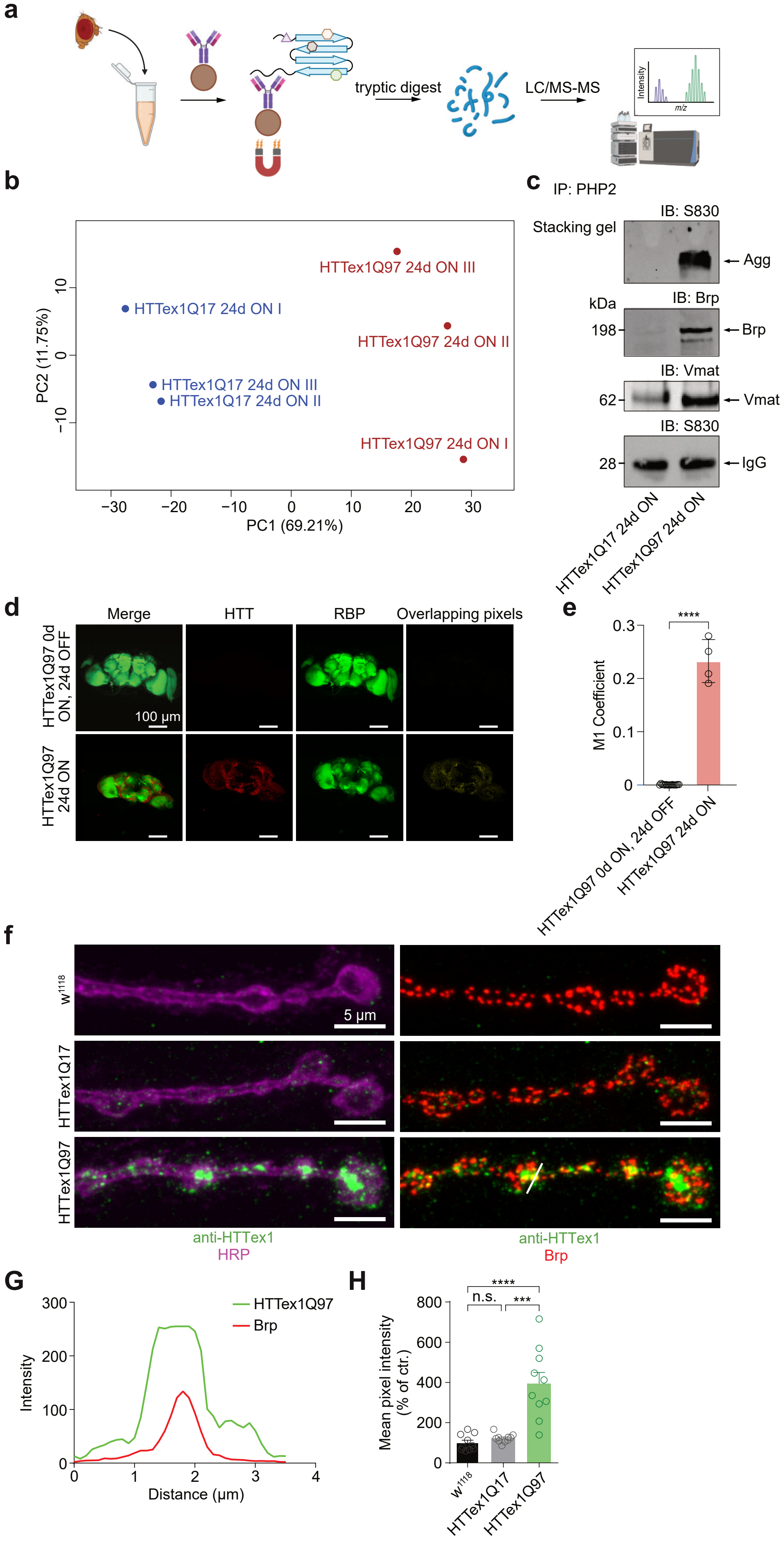


**Supplementary Figure 3**

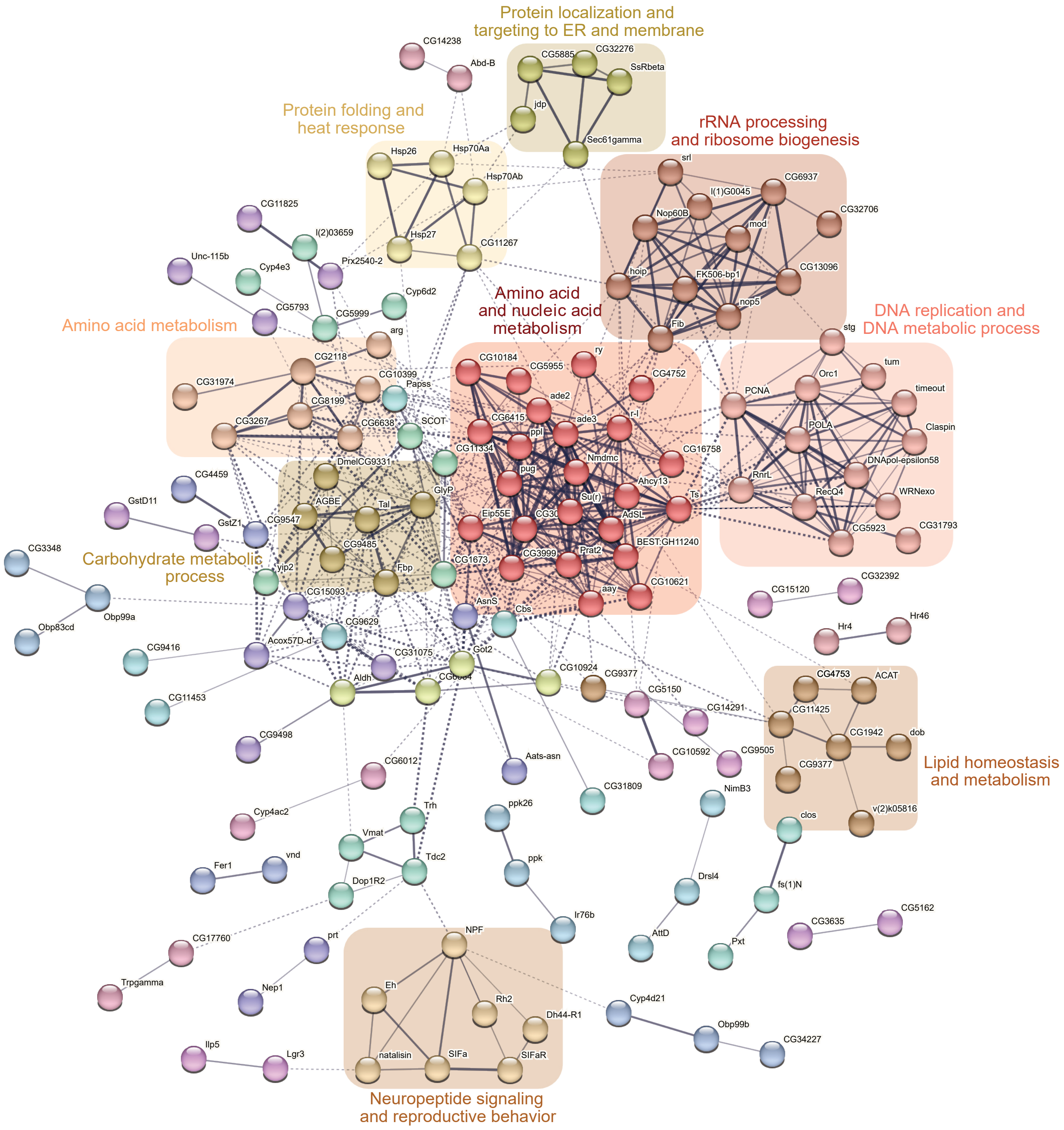


Supplementary Figure 4

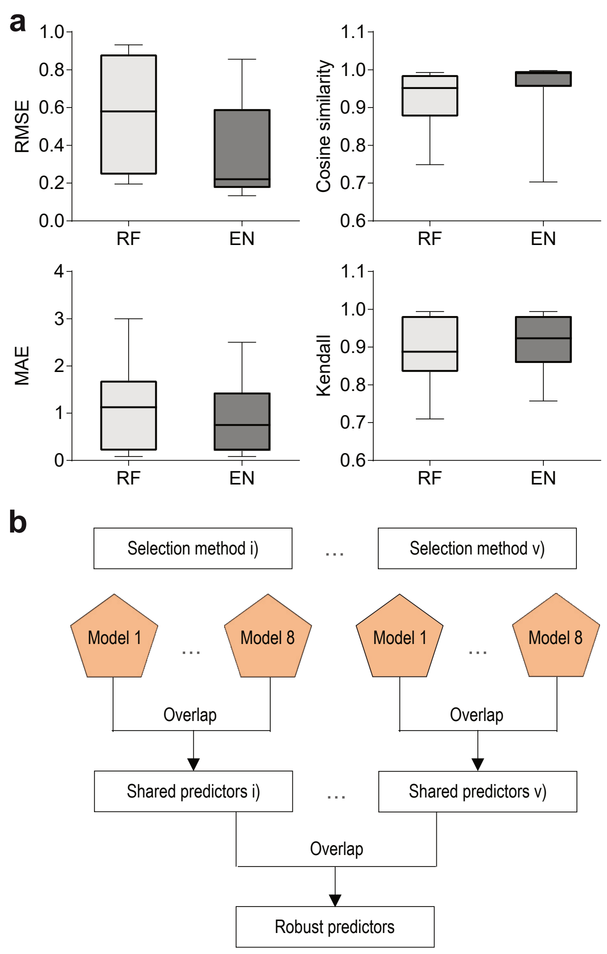


**Supplementary Figure 5**

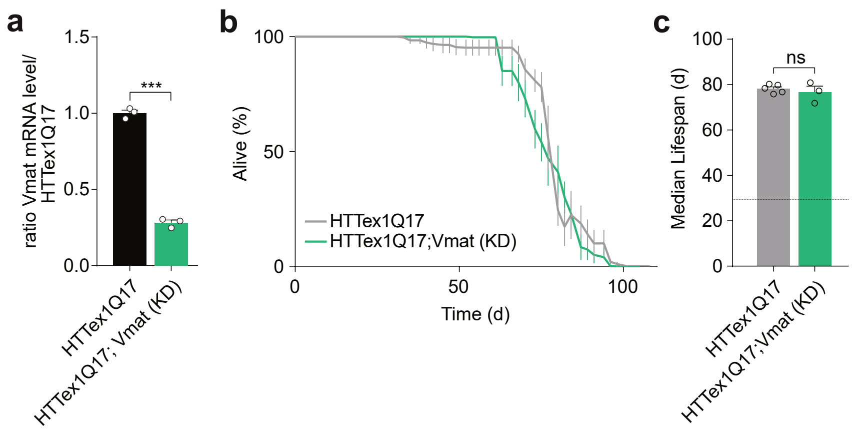


**Supplementary Figure 6**

**Supplementary Figure titles and legends**

**Supplementary Fig. 1: related to Fig. 1: Mutant HTTex1 aggregation in neurons is associated with impairment of locomotor activity**

**(a)** Prediction of the structure of the N-terminal HTT513Q17 protein with a non-pathogenic polyQ tract with AlphaFold3. The predicted protein contains disordered regions and a stably folded $\alpha$-helical N-HEAT domain.

**(b)** Schematic representation of a *Drosophila* activity monitor applied for the quantification of circadian locomotor activity of transgenic HD flies. Laser beams used for fly movement detection are indicated (red).

**(c)** Average locomotor activity of 5-day-old male HD flies pan-neuronally expressing different HTTex1 protein variants with the Elav-GAL4 driver ($n_{HTTex1Q0}$ = 14; $n_{HTTex1Q17}$ = 15; $n_{HTTex1Q49}$ = 15; $n_{HTTex1Q97}$ = 14).

**(d)** Quantification of average locomotor activity of 5-day-old HD flies in **(c)**. Relative activity is displayed as area under the curve normalized to HTTex1Q17. Error bars represent mean ± SEM; one-way ANOVA Dunnett's post hoc test compared to HTTex1Q17; n.s., not statistically significant; *** $p < 0.001$.

**Supplementary Fig. 2: related to Fig. 2: Aggregation of HTTex1 protein variants with and without polyQ interruptions *in vivo* and *in vitro***

**(a)** Quantification of SDS- and heat-stable HTTex1 aggregates formed over time in heads of Elav-GAL4-driven HD transgenic flies by filter retardation assay (FRA). An anti-V5 antibody was applied for aggregate detection.

**(b)** Quantification of SDS- and heat-stable HTTex1 aggregates formed over time in fly heads. Data from FRAs shown in **(a)** were used for quantification.

**(c)** Analysis of motor activity of HD flies pan-neuronally expressing different HTTex1 protein variants with the Elav-GAL4 driver. Motor performance was assessed over time. The number of flies capable to climb a height of 8 cm within 15 sec was quantified. Results are presented as a mean of 3 independent experiments ($n_{Ex1Q75}$ = 100, 98, 98; $n_{Ex1Q61ENK14}$ = 99, 98, 101; $n_{Ex1Q71P4}$ = 95, 100, 98; $n_{Ex1Q17}$ = 100, 100, 92; $n_{w1118}$ = 100, 100, 100 flies).

**(d)** Relative motor activity quantified by calculating the area under the curve from climbing curves in **(c)**. Results are expressed as percentage of control (Elav;$w^{1118}$). Average motor performance quantified in each experiment (n = ~100 flies) is presented as white dots. Bars are the mean ± SEM from 3 independent replicates; statistical significance was assessed by One-way ANOVA Dunnett's post-hoc test; data were compared to Elav;Ex1Q75 transgenic flies.

**(e)** Schematic representation of GST-tagged HTTex1 fusion proteins with and without polyQ interruptions. Orange and blue indicate inserted point mutations in polyQ tracts.

**(f)** Analysis of recombinant GST-tagged (N-term) HTTex1-V5 (C-term) fusion proteins by SDS-PAGE and Coomassie blue staining (left) as well as immunoblotting using anti-GST (middle) or anti-V5 (right) antibodies. (* indicates full-length GST-HTTex1 fusion proteins).

**(g)** Cartoon of a fractionation experiment depicting the generation of pellet and supernatant fractions under non-denaturing conditions by high-speed centrifugation for analysis of spontaneously formed HTTex1 protein assemblies with SDS-PAGE and immunoblotting.

**Supplementary Fig. 3: related to Fig. 3: Accumulation of HTTex1Q97 aggregates in fly neurons is associated with sequestration of synaptic proteins**

**(a)** Simplified co-IP-MS workflow. For co-IPs head lysates prepared from 27-day-old elavGS;HTTex1Q97 or elavGS;HTTex1Q17 transgenic flies after induction of transgene expression with RU486 in 3-day-old flies were incubated with PHP2-coated beads. Tryptic on-bead-digestion was performed before samples were analyzed with LC-MS/MS.

**(b)** Principal component analysis (PCA) of protein abundances of HTTex1Q97 co-immunoprecipitates compared to the HTTex1Q17 co-immunoprecipitates. Samples were enriched by co-IPs from fly head lysates using the PHP2 antibody (see **Supplementary Fig. 3a**). Protein abundances were calculated by employing MaxLFQ-based quantitation.

**(c)** Analysis of PHP2-enriched co-IP samples by SDS-PAGE and immunoblotting (IB). Agg, high molecular weight amyloid-like HTTex1Q97 aggregates in the stacking gel; Brp, Bruchpilot; Vmat, vesicular monoamine transporter; IgG, immunoglobulin light chain.

**(d)** Immunohistochemistry analysis of various brains prepared of elavGS-driven HTTex1Q97-expressing HD fly strains revealed a co-localization of the synaptic RIM-binding protein RBP with HTTex1Q97 aggregates. Images were taken of 27-day-old HD flies after expression of HTTex1Q97 for 24 days (24d-ON) in neurons with RU486.

**(e)** Quantification of the co-localization of RBP with HTTex1Q97 aggregates in inclusion bodies, as observed by immunohistochemistry in fly brains shown in **(d)**.

**(f)** Representative confocal immunofluorescence images of 3$^{rd}$ instar *Drosophila* larvae of the neuromuscular junction (NMJ) expressing Elav-GAL4 labeled for HTT (green) and HRP (horseradish peroxidase, magenta), outlining the NMJ (left panel) and Brp as a presynaptic marker (red, right panel). White line indicates line graph for **(g)**. HTTex1Q97 aggregates at the NMJs expressing Elav;HTTex1Q97 are clearly detectable, while no such structures were detectable in larvae of Elav;HTTex1Q17 or the driver control (Elav;w1118). A partial co-localization of Brp with distinct amyloid-like HTTex1Q97 aggregates at NMJ is detectable; scale bar 5 mm.

**(g)** Line scan analysis supports the co-localization of HTTex1Q97 aggregates with Brp at NMJs.

**(h)** Quantification of the mean pixel intensities of the anti-HTT signal of **(f)** (n = 12, n represents NMJs) supports an accumulation of HTTex1Q97 aggregates in third instar larvae, but not control animals.

**Supplementary Fig. 4: related to Fig. 4: Expression of HTTex1Q97 in young adult neurons through RU486 treatment causes global transcriptome changes in brains of 27-day-old HD flies**

Predicted protein clusters of significantly dysregulated genes detected in 27-day-old brains of RU486-treated elavGS;HTTex1Q97 HD transgenic flies (**Fig. 4a**) but not in RU486 untreated control flies. Only expression changes detected in at least two of the three RU486 treatment conditions (e.g., 3d-ON/OFF and 6d-ON/OFF; see **Fig. 4c**) but not in the RU486-OFF treatment conditions were utilized for protein cluster prediction. Protein clusters were generated with the STRING database using the MCL algorithm. The nine predicted subnetworks are highlighted by an underlying box. Two examples of these predicted transcriptionally dysregulated protein clusters are depicted also in **Fig. 4d** and **e**.

**Supplementary Fig. 5: related to Fig. 5: Machine learning-based analysis of transcriptome and lifespan data reveals genes encoding proteins involved in neurotransmission as positive predictors of fly survival**

**(a)** Performance evaluation of machine learning models: random forests (RF) and elastic net regression (EN) based on four evaluation measures calculated from the comparison of predicted values and ground truth values: root mean squared error (RMSE), cosine similarity, mean absolute error (MAE) of ranks and Kendall's rank correlation coefficient.

**(b)** Scheme illustrating the identification strategy of lifespan predictor genes. The analysis was performed for different sizes of input feature sets: i) no preselection of genes, ii) preselecting the top k=10,000 varying genes from the respective training set, iii) preselection with k=5,000, iv) preselection with k=2,500 and v) preselection with k=1,000. Each time, eight-fold cross-validation across biological groups produced eight models. The intersection of predictive genes from these eight models formed the candidate set of the current preselection and were overlapped with the results from other preselection settings to obtain the final candidates for further investigation.

**Supplementary Fig. 6: related to Fig. 6: Knockdown of Vmat with RNAi in HTTex1Q17-expressing neurons does not significantly influence the survival of elavGS;HTTex1Q17 flies**

**(a)** Quantification of *Vmat* transcript levels in brains of non-RNAi- and RNAi-expressing elavGS-driven HTTex1Q17 transgenic flies. Transgene expression was induced with RU486 in 3-day-old flies and samples were analysed after 6 days. Results of three biological replicates are depicted as white dots and as mean ± SEM; statistical significance was assessed by One-way ANOVA Tukey's post-hoc test.

**(b)** Lifespan analysis of non-RNAi and RNAi-expressing elavGS;HTTex1Q17 flies after induction of transgene expression with RU486 in 3-day-old animals. Lifespan is plotted as the percentage of surviving flies of 5 biological replicates (~50-100 flies each). Results are depicted individually as mean ± SEM of biological replicates.

**(c)** Median life span calculated from survival curves in **(b)**. Average survival of each experiment (n ≥ ~50 flies) is presented as white dots. Bars are mean ± SEM from 5 independent replicates; statistical significance was assessed by One-way ANOVA Dunnett's post-hoc test.

**Supplementary Table titles**

**Supplementary Table 1.** Amino acid sequences of the HTTex1 proteins with and without polyQ interruptions

**Supplementary Table 2a.** Fly proteins significantly co-enriched with mHTTex1 aggregates through co-immunoprecipitation with anti-HTT PHP2 antibody

**Supplementary Table 2b.** Overlapping proteins/genes identified among proteins co-enriched with mHTTex1 aggregates and genes that are differentially downregulated in heads of transgenic flies.

**Supplementary Table 3a.** Transcriptome data sets utilized to define differentially expressed genes (DEGs) in heads of HD and control transgenic flies (for experimental conditions see **Fig. 4a**)

**Supplementary Table 3b.** Genes differentially down- or upregulated in heads of elavGS;HTTex1Q97 vs elavGS;HTTex1Q17 transgenic flies (|LFC| >= 1 and p.adj <0.01); see also **Fig. 4b**

**Supplementary Table 3c.** Genes differentially down- or upregulated in heads of elavGS;HTTex1Q97 vs elavGS;HTTex1Q17 transgenic flies (|LFC| >= 1 and p.adj <0.01); only genes significantly dysregulated in at least two of the three experimental conditions shown in **Fig. 4a** were selected (see **Fig. 4c**)

**Supplementary Table 3d.** Genes differentially down- or upregulated in heads of elavGS;HTTex1Q97 vs elavGS;HTTex1Q17 transgenic flies (p.adj < 0.05); genes were defined without a predefined LFC cutoff (see also **Fig. 4f**)

**Supplementary Table 3e.** GO term enrichment analysis with DEGs (p.adj < 0.05) obtained by comparing transcriptome data of elavGS;HTTex1Q97 and elavGS;HTTex1Q17 transgenic flies (see also **Fig. 4a**)

**Supplementary Table 4.** Gene expression data from elavGS;HTTex1Q97 and elavGS;HTTex1Q17 fly heads collected under different experimental conditions (**Fig. 4a**) for transcriptome analysis

**Supplementary Table 5.** Lifespan measurements of elavGS;HTTex1Q97 and elavGS;HTTex1Q17 transgenic flies propagated under experimental conditions shown in **Fig. 4a**; data were previously reported in Ast et al. 2018

**Supplementary Table 6.** Materials and resources used, including identifiers.
